# Structure-based recasting of a mammalian DNA transpososome as an obligate heterodimer

**DOI:** 10.64898/2026.07.15.738711

**Authors:** Raju Mandal, Alison B. Hickman, Rhea P. Desai, Alexandria Primich, Fred Dyda

**Affiliations:** Laboratory of Molecular Biology, National Institute of Diabetes and Digestive and Kidney Diseases, National Institutes of Health, Bethesda, MD USA

## Abstract

Eukaryotic DNA transpososomes assemble as nucleoprotein complexes containing multiple identical transposase protomers. We determined the structure of the hyperactive *Myotis lucifugus* piggyBat transpososome and discovered that it uses an unusual crescent-shaped, asymmetric tetramer to synapse divergent inverted terminal repeats. We found that identical amino-acid sequence motifs adopt distinct roles to mediate two modes of DNA binding: one to perform strand transfer and one, devoid of catalytic activity, that promotes synapsis while simultaneously protecting the transposon from auto-destructive internal cleavage by its active sites. Guided by the observed modularity of the assembly, we engineered an obligate heterodimeric system by identifying mutations that suppress homodimer formation and paired this with specific point mutations that prevent non-targeted integration. By adding to the heterodimer two different TALE domains designed to bind a human genomic safe harbor sequence, we achieved >98% targeted integration at the intended sequence in a plasmid-based assay, validating the viability of heterodimeric transposases for genomic applications.

## Introduction

The integration of therapeutic genes into the human genome is a critical challenge for gene and cell therapy.^1–3^ Transposons - and the *piggyBac* superfamily of DNA transposons in particular - have emerged as promising tools for these applications due to their ability to integrate large DNA cargos into both dividing and non-dividing cells.^4,5^ However, a major safety concern for all cut-and-paste DNA transposons including *piggyBac* is their random integration into the host genome. This can lead to gene disruptions and has resulted in adverse clinical outcomes. In principle, this safety risk could be mitigated if transposon integration could be directed to specific, “safe” locations in the genome, known as Genomic Safe Harbors (GSHs).^6^

Cut-and-paste transposition proceeds by an assembly of the transposon-encoded transposase protein that, through its inherent nuclease activity, is able to capture, synapse and process the Left End and Right Ends (LE and RE) of the transposon. DNA motifs that are recognized by various parts of the transposase are often arranged as imperfect Terminal Inverted Repeats (TIR) at the two transposon ends. Cut-and-paste transposition can be broadly divided into two mechanistic phases, excision and integration (Figure 1A). Excision liberates the transposon from the donor site, and generates staggered Double Strand Breaks (DSB) on flanking DNA. Unlike many other transposons, members of the *piggyBac* superfamily generate a hairpin intermediate at each transposon end upon excision. As *piggyBac* transposons always target a TTAA tetranucleotide for integration (which becomes the characteristic target site duplication (TSD) when integration is complete), TTAA always flanks the transposon, as the donor site is the product of the last round of integration. Due to the staggered cuts made by the transposase at the transposon ends, the donor flanks that remain have short, complementary overhangs and are sealed seamlessly by the cellular DNA repair machinery. These seamless joints and therefore the lack of genomic scars (or “excision footprints”) are another key characteristic of the *piggyBac* superfamily.

**Figure 1.**
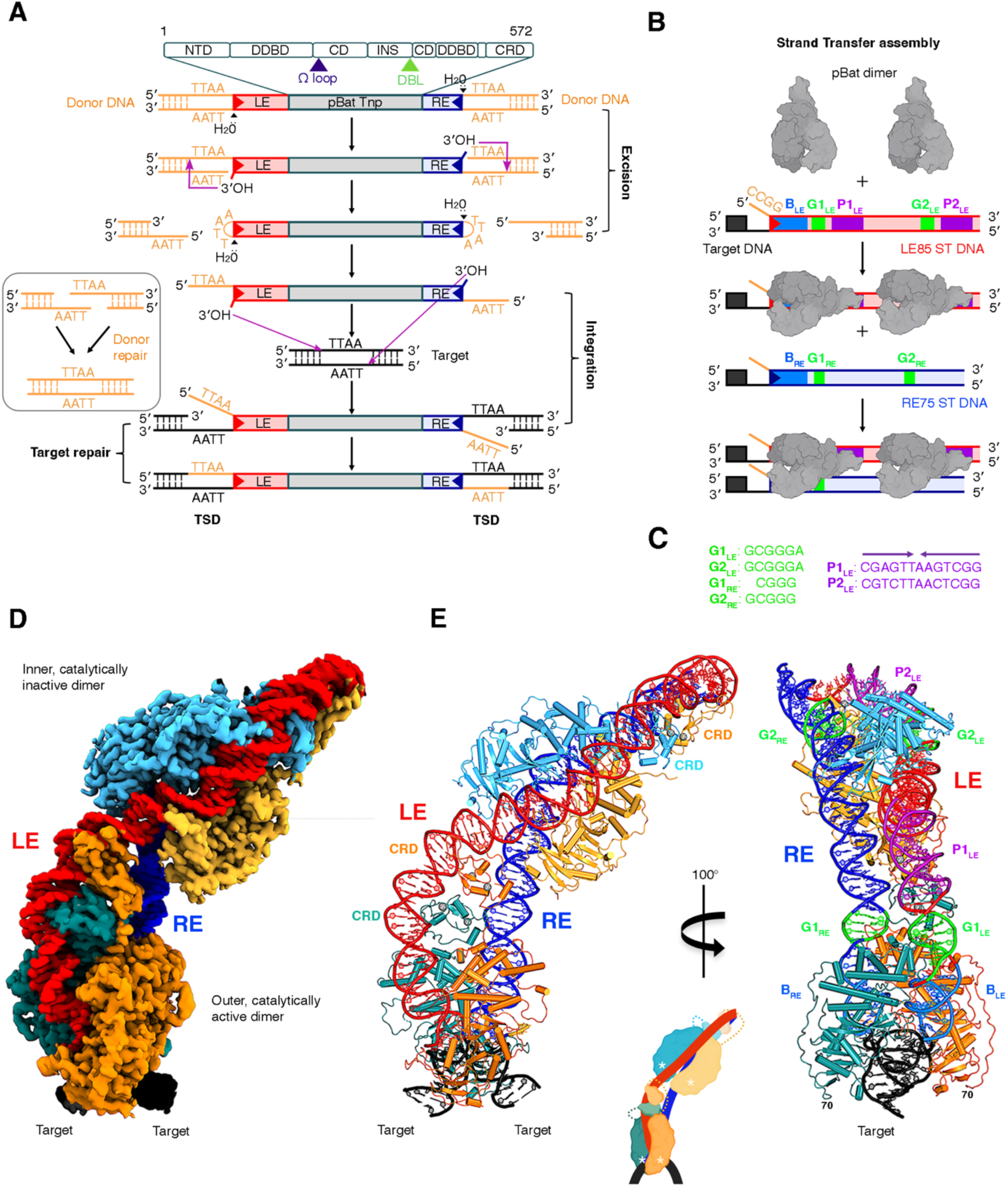
Cryo-EM structure of piggyBat transpososome reveals its crescent-shaped architecture. (A) Schematic showing the transposition pathway of *piggyBac*-like genetic elements. The domain structure of the piggyBat (pBat) transposase is shown on top (NTD, N-terminal domain; DDBD, dimerization and DNA-binding domain; CD, catalytic domain; INS, insertion domain; CRD, cysteine-rich domain). The locations in the primary sequence of the Ω-loop (purple triangle) and Differential Binding Loop (DBL, green triangle) are indicated. (B) Schematic overview of the preparation of the strand transfer complex for cryo-EM analysis from purified pBat transposase, LE85, and RE75. Recurring binding motifs on the LE and RE are shown in blue, green, and purple. (C) Aligned sequences of the “G” motifs present on both transposon ends (shown schematically in green in (B)) and the palindromic “P” motifs present only on the LE (shown schematically in purple in (B)). (D) Cryo-EM density of the composite DeepEMhancer-sharpened map of the pBat STC complex contoured at 0.013. Four identical protomers, assembled as two dimers, synapse the LE and RE. (E) Atomic model of the STC complex, viewed in two different orientations. On the right side, the binding motifs on the LE and RE are colored as in (B). Inset shows a schematic of the assembly, with active sites marked with white asterisks.

Nucleoprotein complexes containing the two liberated TIRs then proceed to capture the target DNA. At this point, the key intermediate is the transpososome that contains the TIRs and target DNA, and it catalyzes the two strand-transfer events by which the transposon’s transferred strand becomes covalently attached to the target DNA.

Efforts to achieve targeted transposition by fusing transposases to site-specific DNA-binding proteins—such as zinc fingers (ZnF), transcription activator-like effectors (TALEs), or dCas9—have met with limited success.^7–9^ A fundamental problem with these fusions lies in the architecture of the transpososomes themselves that can contain homodimers, homotetramers, or even larger homomeric complexes of the transposases. Fusing a targeting domain to a transposase in this context means multiple, identical targeting domains are present, yet target binding needs only one of these to recognize the intended target sequence. Efficient targeting of symmetrical target sites is possible with identical targeting domains, but this is a significant limitation. A second problem is that transposases themselves have inherent non-specific target DNA binding function, without which they would be unable to integrate. This activity, unless suppressed, will direct integration to non-specific sites independent of the presence of a fused specific binder, giving rise to frequent and undesired “off-site” integrations. Progress to the rational design of more specific systems has been hindered by a lack of understanding of the precise three-dimensional structure of active transpososomes of the most relevant eukaryotic DNA transposons such as *piggyBac* and *Sleeping Beauty*.

Our work addresses this knowledge gap using the *piggyBat* (*pBat*) transposon from the bat *Myotis lucifugus*, the only known active DNA transposon in a mammalian genome.^10,11^ Although wild-type *pBat* has modest activity, we have recently showed that its transposition activity can be substantially enhanced through modifications.^12^ For example, *pBat* possesses three binding sites for the *pBat* transposase (pBat) dimer at its transposon LE, but removing the innermost of these markedly stimulated activity. Removing yet another binding site, however, led to substantial loss of transposition activity suggesting that maximal activity requires two binding sites and thus a tetrameric assembly.^12^

Our biochemical data^12^ also revealed that the pBat/DNA binding affinities to the LE and RE are substantially different and that transposons with symmetrized ends (LE+LE or RE+RE) are inactive. Here, we exploited this difference in affinity to assemble a functional and hyperactive transpososome composed of two pBat dimers and oligonucleotiodes representing LE andRE sequences that maximizes transposition activity in cells. The oligonucleotide design also includes target DNA enabling us to capture the integration intermediate. Using single-particle cryo-electron microscopy (cryo-EM), we have resolved the three-dimensional structure of this crucial active state of the *pBat* transpososome linked to its target DNA at 2.9 Å resolution. The structure shows the architecture of the complex that synapses two very different transposon ends without the need for host factors or any other transposon-encoded proteins. The structural information allowed us to redesign the *pBat* transpososome into a functional heterodimer, providing a foundational step toward the rational design of targeted, safe, and efficient gene editing tools.

## Results

### The architecture of *pBat* transpososome complex is crescent-shaped

To determine the cryo-EM structure of the active tetrameric *pBat* transpososome, we staged the assembly of the integration or “strand transfer” complex (STC) in which the two transferred strands of the transposon are covalently attached to target DNA. The STC is on the transposition pathway (Figure 1A), and represents the crucial step of capturing the target at the obligatory TTAA tetranucleotide, a unique characteristic of the *piggyBac* superfamily. We took advantage of the high affinity of the *pBat* transposase to its LE, and first formed complexes of two transposase dimers bound to one oligonucleotide consisting of 85 bp LE in which the transferred strand at the transposon end was linked covalently to 15 bp of target DNA (Figure 1B). These complexes were chromatographically isolated and then we added a 75 bp long RE, which was also extended covalently by 15 bp of target DNA. The resulting complexes therefore contained a tetramer of the transposase bound to LE85, RE75 and the target DNA after strand transfer (Figure S1). They resulted in good quality particles (Figures S2A and S2B), and 2D classifications were dominated by crescent-shaped projections of the tetrameric species (Figure S2B).

The final electrostatic potential density maps (Figures S2C, S3, and S4) after reference-based motion correction had sufficient resolution to build both the protein and nucleic acid components manually and unambiguously. The final map obtained after focused refinement of the individual dimers and associated nucleic acid improved the resolution beyond 3 Å in many parts of the assembly (Figures S4E-J). The transpososome shows the two dimers synapsing one LE and one RE (Figures 1D and 1E). Consistent with the 2D class averages, the assembly has an unusual bent shape, rather than the typical approximate low-order rotational symmetry seen in other DNA transpososomes resolved to date. This shape is however consistent with the asymmetric distribution of binding motifs on LE and RE (labelled “B”, “G”, and “P” in Figure 1E, right) that occur at different distances from the tips of the transposon where the target DNA is attached.

Remarkably, the assembly shows no evidence for protein-protein interactions between the two dimers (Figures 1D and 1E). In the outer, catalytically active dimer, the DDD residues that constitute the two active sites (D237, D309, D413) coordinate to the scissile phosphate of the transferred strand through bound Ca^2+^ ions that do not support strand cleavage or transfer (Figure 2A, inset bottom), thereby trapping the assembly state. In contrast, the inner dimers are catalytically inactive and the phosphoribose backbone is comfortably away from both the active sites (Figure 2A, inset top). Thus, the two dimers synapsing LE and RE have different binding modes despite the similar DNA sequences involved and the obviously identical protein sequences. A key factor driving this asymmetry is the large bend of the LE introduced by the two CRDs of the outer dimer that dimerize on the LE outer purple palindrome (Figures 1C and 2B) such that the trajectories of the DNA going into the inner dimer from the outer one is different from those of the DNA entering the outer dimer at the tip of the transposon (compare Figures 2C and 2D). Therefore, the two dimers, in the absence of protein-protein contacts, appear to be communicating their DNA-bound state to each other via the enforced DNA configuration.

**Figure 2.**
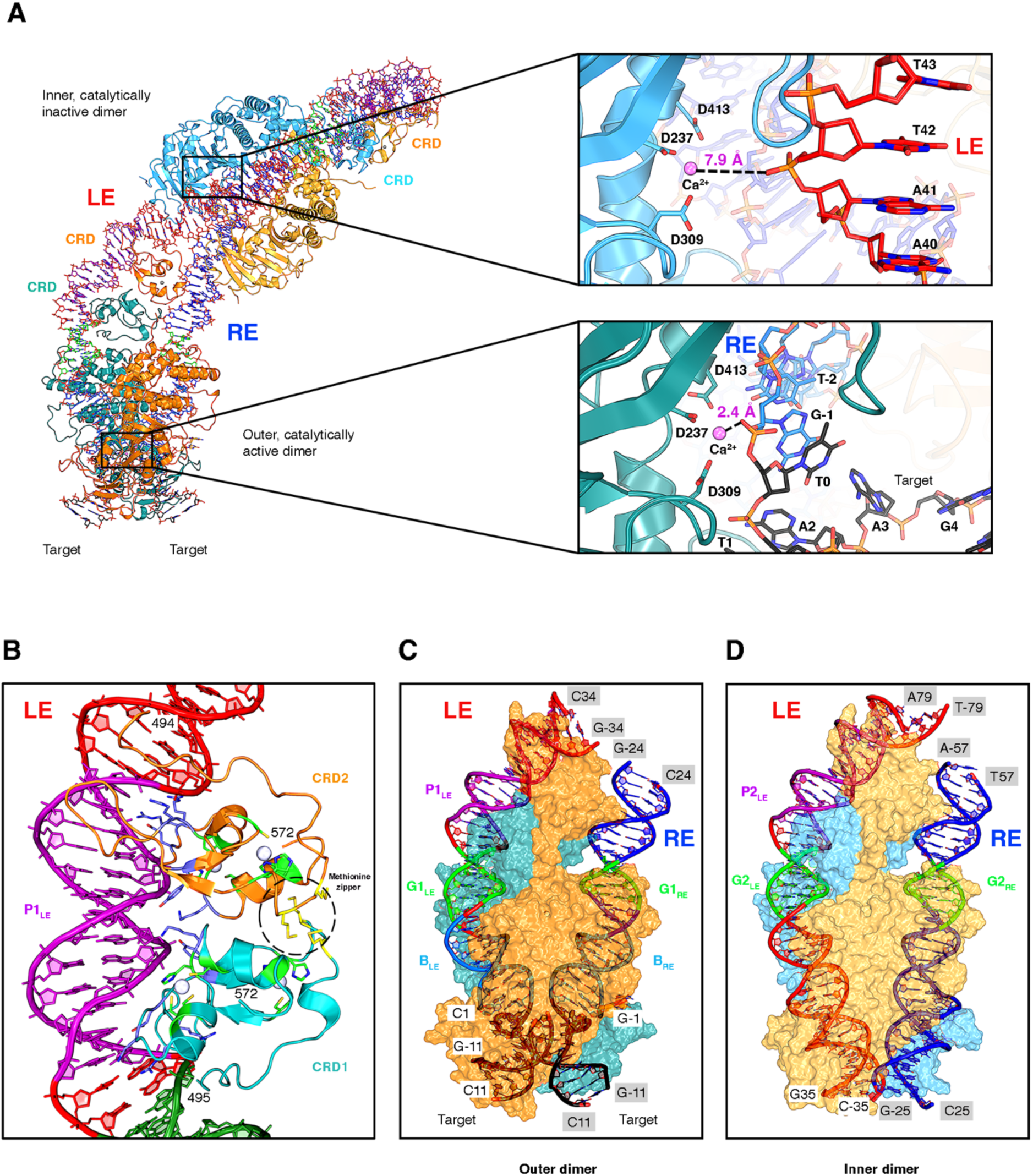
Outer catalytic dimer and inner inactive dimer have different active site configurations and DNA end trajectories. (A) Overall view of the tetramer and close-ups of active sites. (Top) Active site of one monomer in the inner dimer, showing the metal-ion coordinating residues and the closest phosphate to the Ca^2+^ ion. (Bottom) Active site of one monomer in the outer active site, showing metal-ion coordinating ligands and the scissile phosphate at the transposon RE TIR. B) Detailed interaction between the CRD domains and the purple box region of LE DNA. The interacting residues are highlighted in sticks, and the zinc metal ions are shown as gray spheres. (C) Trajectories of LE bp 1-34 and RE bp 1-24 through the outer dimer. (D) Trajectories of LE bp 35-79 and RE 25-57 through the inner dimer.

The LE bends more than 90° centered around bp 25 as it runs through the outer dimer and bends again by about 90° again around bp 69 due to the binding of the CRDs contributed by both dimers (Figure 1E). The second bend is roughly in a plane that is perpendicular to the first bend. These LE DNA distortions are much greater than the ∼40° and ∼60° LE DNA bends that were observed in previous pBac or pBat structures,^12,13^ suggesting that the final active conformation of the LE requires the RE and the DNA-bound inner dimer. In contrast to the sharp bends of the LE, the RE is essentially straight as it runs through the outer dimer and bends only slightly in the inner dimer, with only very minimal interactions with the CRDs of both outer and inner dimers. Taken collectively, the very different paths of LE and RE DNAs is a striking feature of the transpososome assembly, and reflect the different binding motif distributions on the LE and on the RE including the lack of CRD binding sites on the RE.

Another contributor to the geometric asymmetry of the assembly is the asymmetry in how the *pBat* transposase domains interact with LE and RE. In particular, one transposase monomer forms protein/protein interactions with its own CRD (Figures 1D and 1E and inset; for example, the teal monomer in the outer dimer) which in turn dimerizes with another CRD (orange, Figure 1D and 1E) through a methionine zipper (Figure 2B). This arrangement requires a linker between the transposase core and the CRD that is long and flexible enough to cover two different distances in space, 17 Å and 38 Å, between the two cores and the two CRDs. The linker (approximately residues 480-494, depending on which monomer) is not visible in the potential density, perhaps not unexpectedly given the two different conformations it has to assume (Figure S5). Nevertheless, the geometry of the arrangement is such that the monomer whose active site is processing the RE transferred strand is connected to the CRD proximal to the transposase core, while the other monomer is connected to the distal CRD (see Figure 1E and inset). The alternative connectivity between cores and CRDs appears sterically impossible as the linkers would have to loop around both protein and DNA. Another consequence of the asymmetric binding mode in which the CRDs interact only with the LE is that there are about one and half turns of naked, solvent-exposed dsDNA (bps 27-40) of RE between the outer and inner dimers of the STC assembly. The details of the interactions between pBat and both DNAs are shown in Figures S6 and S7.

### The Omega loop and Differential Binding Loop act as guardians of the transposon

Given the experimental observations that optimal transposition activity is observed with the tetramer, and the dimer alone has minimal activity, a key mechanistic problem that pBat has to solve is how to licence nuclease activity at the transposons tips while preventing it inside of the transposon where the inner dimer is bound. Wanton nuclease activity by the inner dimer would damage the transposon and likely result in the loss of mobilizability and/or the loss of remobilizability as crucial transposase binding motifs closer to the tip would be lost. Guarding the transposon DNA is achieved by two sequence segments of the transposase that adopt different configurations in the inner and outer dimers, resulting in significant differences in how they interact with the DNA in the two dimers.

The Ω loop (residues P242-G261) is an inserted motif in the RNaseH-like catalytic domain and is a highly conserved segment of the *piggyBac* superfamily. In the catalytically active outer dimer, the two loops play a crucial role in recognizing the TTAA target motif (Figure 3A, left). In contrast, in the inner dimer, the two Ω loops assume distinct conformations in each monomer and different modes of interaction with LE and RE, and engage mainly with the phosphoribose backbone (Figure 3A, middle and right). Taken together, we observed three significantly different Ω loop conformations, and three different modes of interactions with the DNA by this 21 amino acid long highly conserved motif.

**Figure 3.**
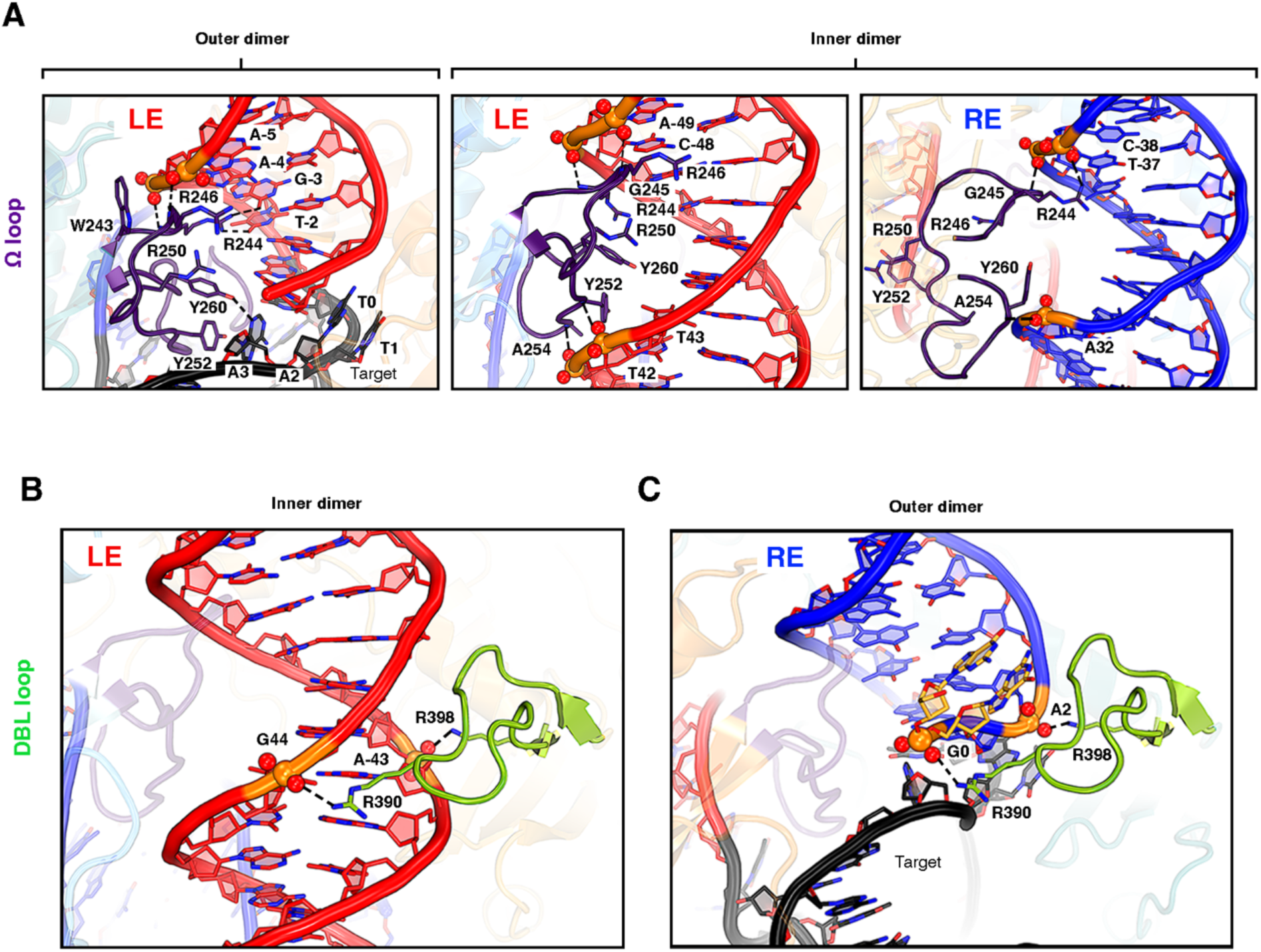
Comparison of the omega (W) loop and the differential binding loop (DBL) conformations in the inner and outer dimers. (A) Close-ups showing the three different conformations of the W loop (amino acids 242-261, shown in purple). In the outer dimer, only the LE interaction is shown as it is very similar to that on the RE. In contrast, in the inner dimer, two very different conformations and sets of interactions are observed on the LE and the RE. Backbone phosphate groups contacted by omega loop residues are shown in ball-and-stick representation (P, orange; O, red). (B) Close-up showing the different conformations of the DBL (shown in green) in the inner dimer and (C) outer dimer.

The second structural element responsible for the distinct DNA binding modes of the inner and outer dimers is a segment between M382-P399 that forms a twisted β hairpin loop. This “Differential Binding Loop” (DBL) has R390 at its tip that dives deeply into the minor groove of both LE and RE at the inner dimers (Figure 3B) and narrows the grooves; it has only cursory interactions with the non-transferred strand of the outer dimers (Figure 3C). Interestingly, a recent extensive metagenomic analysis of the *piggyBac* superfamily identified the DBL as a particular feature of the group of the superfamily to which *pBat* belongs.^14^ As we have previously shown, this group is also distinguished by a CRD domain topology that differs from that of the *Trichoplusia ni piggyBac* transposase (pBac).^12^

Together, the Ω loop and the DBL are largely responsible for the distinct DNA binding modes of the outer and inner dimers. The CRD dimer of the outer dimer introduce a large bend but only in the LE. This results in the transposon DNA entering the catalytic domains of the inner dimer with trajectories such that the engagement of the Ω loops of the inner dimer, together with the interactions provided by the DBLs, can keep the phosphoribose backbones safely away from the inner dimer active sites. All of these structural elements combine to ensure the integrity of the transposon DNA and protect it from undesirable and transposon-debilitating cleavage.

### The structure allows the identification of integration-deficient point mutants

Mutants that are defective for integration but retain precise excision (Exc+Int−) have been identified for a number of cut-and-paste DNA transposases.^15,16^ These mutants have been shown to be of great utility, for instance to remove an integrated transposon from a genome while preventing reintegration. They are also potentially useful for site-specific targeting applications in circumstances where a fused site-specific targeting domain could provide the necessary target capture activity without relying on the non-site-specific target capture activity of the transposase. In principle, this approach could suppress unintended non-targeted random insertions, one of the main limitations of traditional transposon-based site-specifc genomic applications, as integrations should occur only at the intended target site. The key is to identify amino acids that are important for non-specific target DNA binding, but do not have a role in the chemical steps of transposition or in the stabilization of necessary conformational intermediates. For transposases from the *piggyBac* superfamily, it is also important to exclude those amino acids that contribute to the recognition of the TTAA motif that is essential for both for excision and for integration.

We used the STC structure to identify candidate Exc+Int− mutants in pBat, by focusing on those amino acids that interact only with target DNA. Four amino acids (Y281, K256, R333, and R336) were mutated individually to alanine and the impact on excision and integration activities assessed (Figure 4A). (We also included E238A, as our interest was piqued by its location very near the enzyme active site.) HEK293T cells were transfected with donor and helper plasmids and excision activity was measured after 24 h and 48 h, as determined by PCR detection of the repaired donor plasmid (Figure 4B). Transposition activity was measured by colony count assays following puromycin selection initiated 48 hr after transfection (Figure 4C). The two mutants with the lowest excision activity (Y281A and K256A) had no detectable transposition activity, whereas R333A exhibited substantial residual transposition activity. On the other hand, R336A maintained robust excision activity 24 h after transfection and had low residual transposition activity, so we chose to focus on R336A for further work. Unexpectedly, we found that the E238A mutant behaved as a catalytic mutant, completely abolishing excision.

**Figure 4.**
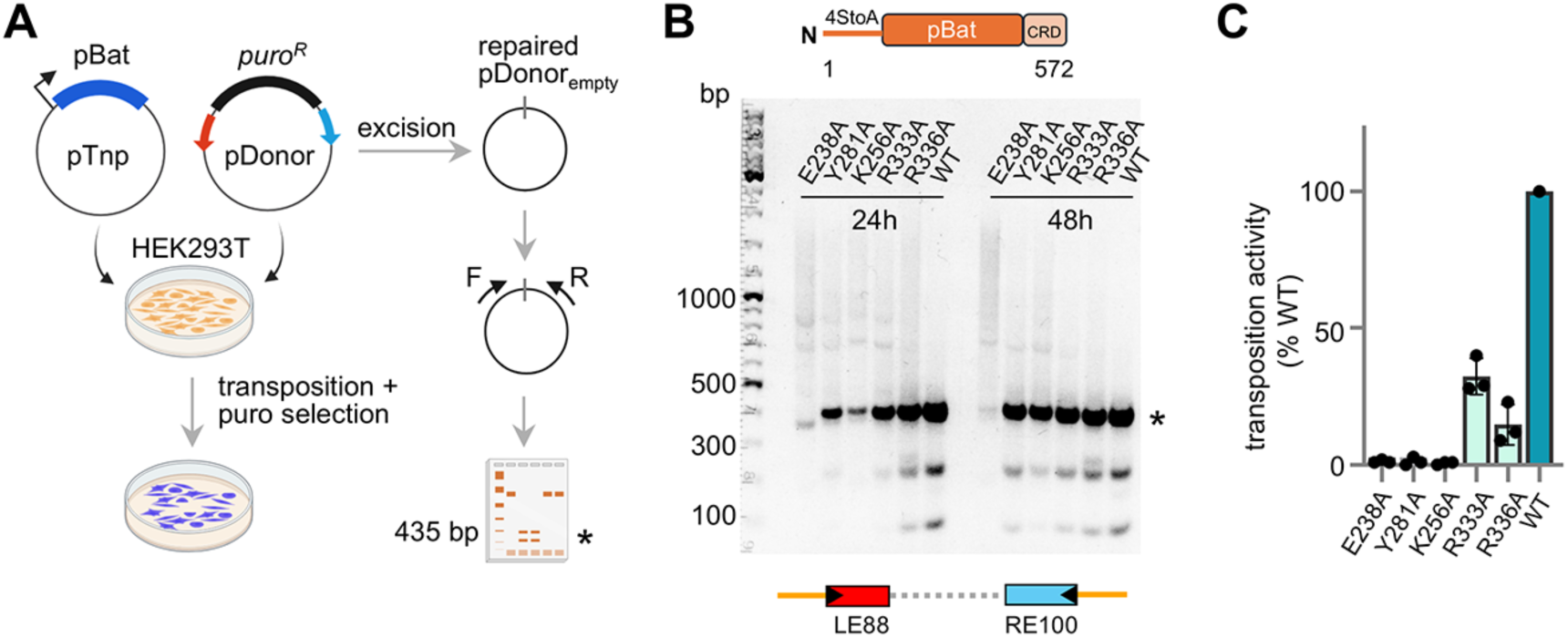
The effect of target-contacting point mutations on transposon excision and integration. (A) Schematic of the two-plasmid assay in HEK293T cells to detect excision and integration. Figure prepared using BioRender. (B) Agarose gel showing the effect of point mutations on excision as detected by the PCR product (marked with black star) corresponding to the repaired pDonor joint. The assay was done using 4StoA pBat transposase (designated “WT” here) and LE88-RE100 transposon, and was repeated n=3. (C) Results of colony count assay for integration for point mutants relative to 4StoA pBat.

### The pBat dimer interface is unaffected by transposon end binding

Both *pBac* and *pBat* transposases form dimers prior to DNA binding when expressed in mammalian cells.^12,13^ This apparently obligatory dimerization is not dependent of the CRD domain,^17^ indicating that domains upstream of the CRD are responsible. Although the observed dimer interface of the pBat dimer when bound to DNA is larger than that of pBac, 1103 Å^2^ vs only 653 Å^2^, it is still relatively modest in size. However, TIR binding by pBat involves a large number of interactions with both of the pBat cores (resides W71–D476) of the dimer, leaving the question open of whether the same interface stabilizes the apo-dimer in the absence of DNA. In particular, we wondered whether there might be a role for the N-terminus sequence upstream of residue 70, the first N-terminal residue visible in the potential density maps of the transpososome, prior to TIR binding.

To address the question of the nature of the pBat dimer interface prior to DNA binding, we obtained a modest resolution cryo-EM structure of DNA-free pBat residues 1-498 (Figure 5A). As we have not succeeded in expressing soluble N-terminally truncated versions of pBat, this construct was used as it forms dimers in solution and we wanted to avoid any ambiguity involving the possible dimerization of the CTDs (Figure S8). Although a preferred orientation problem limited the quality of the reconstruction so that an atomic model could not be built with confidence (Figures S9 and S10), the map was clear enough to show density features at the monomer-monomer interface that did not differ from those of the dimer interface seen in the STC structure. Furthermore, we could not assign any potential density to residues upstream of amino acid 70, suggesting that they are not playing a crucial role in dimerization. We conclude that the observed 1103 Å^2^ interface is sufficient for the obligatory dimerization of the *pBat* transposase.

**Figure 5.**
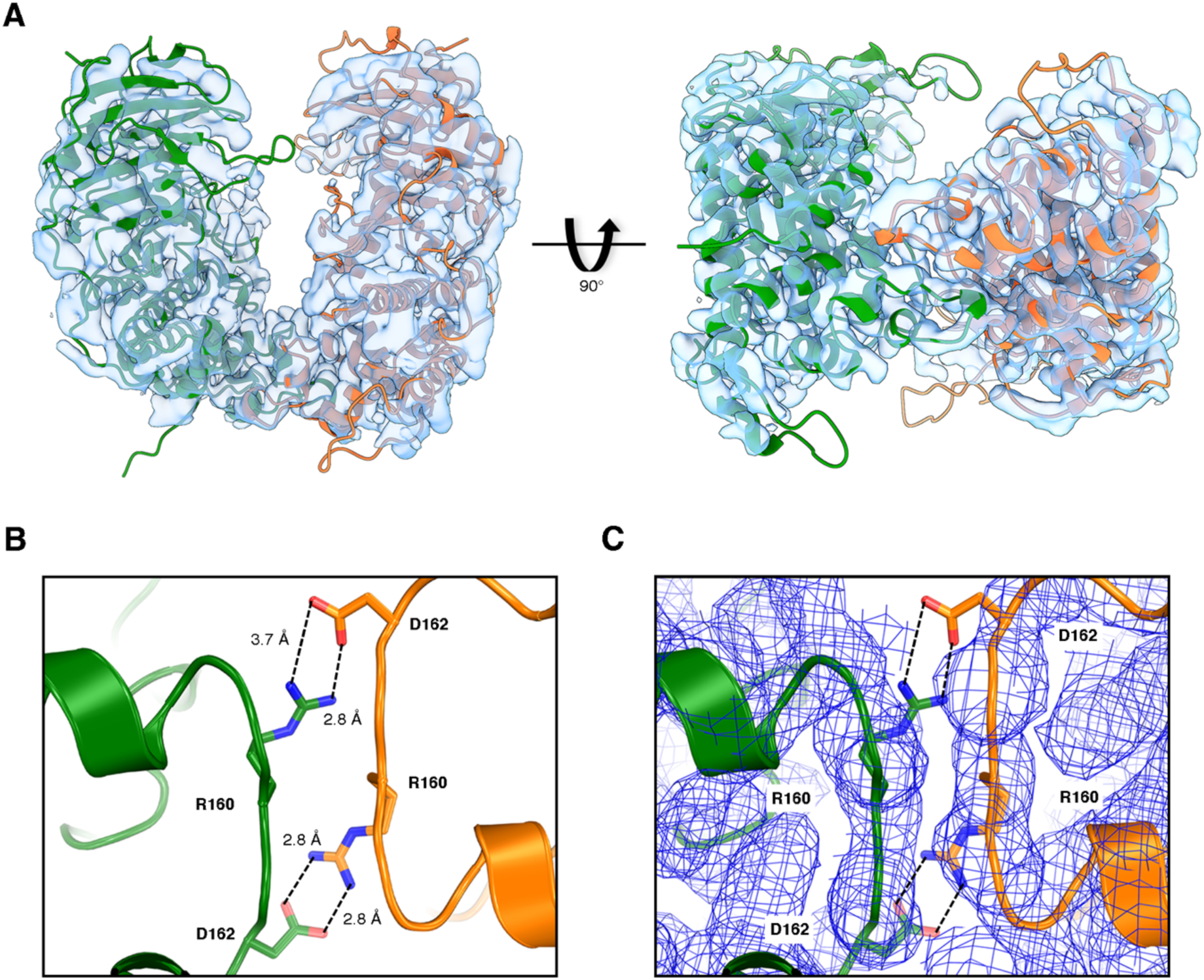
The dimeric interface. (A) The fit of apo pBat transposases into the cryo-EM density map, shown in two different orientations. The dimeric interface is highlighted on the right. (B) Representation of ion pairs present in the dimeric interface. Their direct interactions provide stability to the pBat dimer. (C) Ion pair residues and their fit into the low-resolution cryo-EM density map. The sharpened map was contoured at 3.5 sigma, shown in blue.

To gain more insight into the structural basis of pBat dimerization, we calculated the electrostatic isosurfaces with APBS 1.4^18^ both for the core domain and for a monomer of the core domain; these positive isosurfaces at 11 kT/e are shown in Figures 6A and 6B. In the case of the dimer, the isosurface extends far into solvent, but much less so for the monomer, suggesting that core dimerization promotes the capture of the TIRs. Furthermore, binding of the subterminal inverted repeats by the CRDs (motifs P1_LE_ and P2_LE_, Figure 2B) likely require CRD dimerization to bend the DNA, and as each monomer supplies one CRD, core dimerization is needed to make two CRDs available. Taken together, the data suggests that dimerization is required for TIR binding.

**Figure 6.**
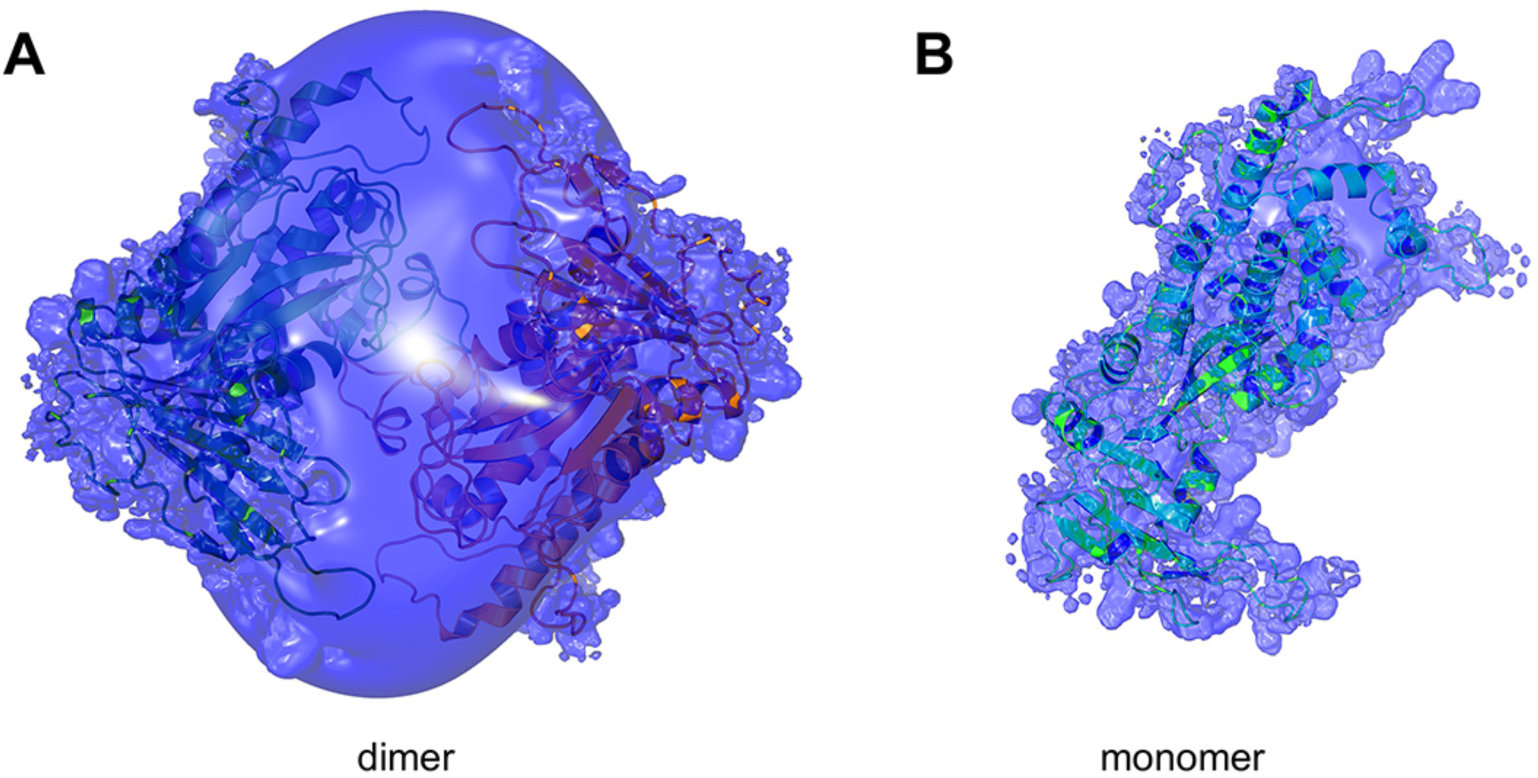
Electrostatic comparison of piggyBat transposase dimer and monomer. Positive electrostatic isosurfaces at 11 kT/e for the core dimer (A) and for the isolated core monomer (B). The calculation was performed with APBS 1.4, Amber charges were assigned using PDB2PQR.^47^

As seen in the STC complex, the N-terminal amino acids upstream of residues 70 are also disordered in the apo form of the transposase, consistent with the results from the flDPnn2 disorder prediction tool (Figure S5).^19^ As has been observed across the *piggyBac* superfamily, this region includes a number of negatively charged residues. Residues 1-69 of pBat has an estimated^20^ pI ∼4.1 in addition to several potential Casein Kinase II phosphorylation sites. In contrast, the folded core domain dimer has a calculated^21^ pI of 10.0 when the N-terminus is excluded. In light of this, we cannot rule out the possibility that the disordered N-terminus contributes to dimerization by stabilizing the pBat dimer through charge compensation prior to TIR binding. Unusually, core dimerization appears to be dependent on a number of polar interactions in the interface, involving buried ion pairs formed by residues R160 and D162 across the interface (Figures 5B and 5C) and only a modest reliance on hydrophobic interactions.

### pBat can be re-engineered as an active, heterodimeric DNA transposase

Encouraged by features of the observed dimer interface and TIR binding, we sought to redesign the *pBat* system from an obligatory tetramer into one in which a heterodimer would be sufficient to carry out transposition in mammalian cells. We reasoned that an obligatory heterodimer could serve as the platform for improved genomic targeting of integration as two different targeting domains, designed to bind to sequences flanking a specific TTAA tetranucleotide, could be fused to each of the monomers. We further hypothesized that the inner dimer is needed only to contribute DNA binding affinity for productive synapse of LE and RE, and impacts neither the chemical steps of transposition nor the ability of the outer dimer to capture target DNA. If so, as the CRDs are major contributors to TIR binding, asymmetry and sufficient binding affinity might be achieved by linking two pBat CRDs to the C-terminus of one pBat core at the same time as linking a different site-specific DNA binding domain to the other. Redesigning the RE to include the binding site for the new site-specific DNA binding domain should allow the assembly of functional heterodimers (shown schematically in Figure 7A).

**Figure 7.**
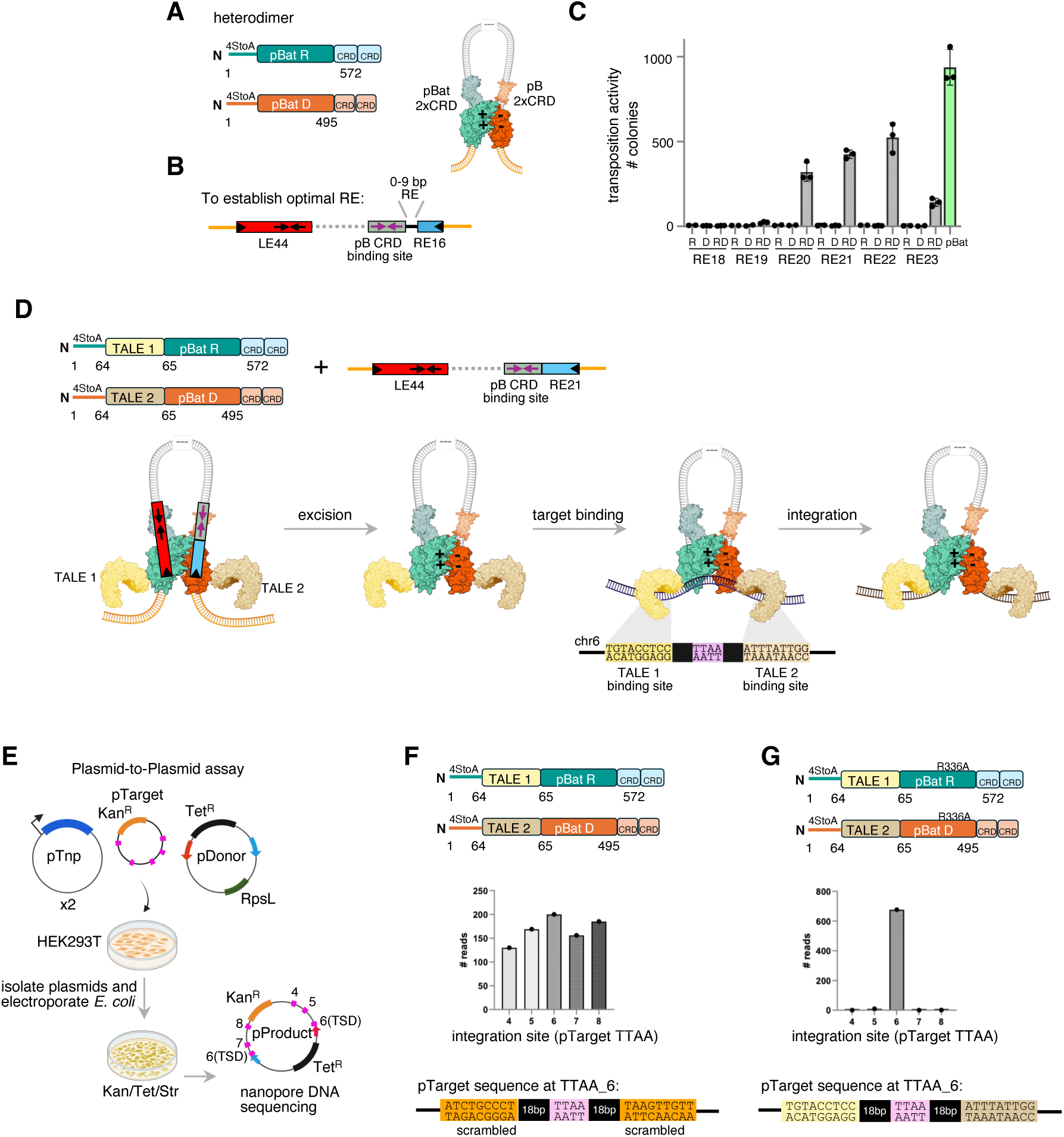
Design strategy and characterization of an active obligate heterodimeric pBat transposase. (A) Schematic of design of two modified pBat transposase monomers and their proposed assembly on modified transposon ends. One monomer (orange) consists of the pBat transposase (residues 1-572) with an additional CRD domain appended to its C-terminus, the other (teal) is the pBat core linked to two tandem pB CRDs. “R” and “D” indicate mutations in the core interface, D162R and R160D, respectively and are represented as “+” and “-” to highlight the complementary ion pairs. (B) Schematic of transposon library used to establish best spacing between the pBat RE tip and the binding site for the two tandem pB CRD (C) Transposition activity as assessed by colony count assay for the transposon variants depicted in (B). “R” and “D” indicate the activity when each modified transposase was assayed alone, “RD” indicates the activity when they were assayed together. For comparison, the activity of the hyperactive pBat system, 4StoA pBat + LE88-RE100, was included (far right, in green). The assay was repeated more than three times, with similar results. (D) Schematic showing the experimental design for the TALE-transposase heterodimer, and the proposed pathway. The TALE 1 binding site is located 18 bp upstream of the targeted TTAA, the TALE 2 binding site 18 bp downstream. Figure created using Biorender. (E) Schematic of the plasmid-to-plasmid assay used to assay targeting. Figure created using Biorender. (F) Distribution of integration events into the five TTAA sites in pTarget for the TALE-heterodimer combination when the TALE binding site sequences around site TTAA_6 are scrambled. The assay was repeated more than three times, with similar results. (G) Distribution of integration events into the five TTAA sites in pTarget for the TALE-heterodimer combination when the TALE binding site sequences around site TTAA_6 are intact. The assay was repeated more than three times, with similar results.

To test this hypothesis, we generated two modified *pBat* transposases: pBat-2xCRD to bind the LE, and pBat-pB2xCRD for RE binding in which two pBac CRDs were appended to the pBat 1-495 core, as their binding mode is known^13^ and they recognize a palindromic sequence unrelated to that of pBat. In these new transposase variants, we also mutated the buried reciprocal ion pairs formed by R160 and D162 in the core domain dimer interface. Thus, one transposase contains the WT R160 and D162R mutation (“R”, teal in Figure 7) and the other contains the R160D mutation and WT D162 (“D”, orange in Figure 7). We theorized that electrostatic charge incompatibility would inhibit the formation of homodimers of either of the two new *pBat* transposase constructs when expressed in cells.

To test the heterodimeric transposase concept in HEK293T cells, we generated a series of pDonor plasmids that contained minimal transposon ends (Figure 7B). On the LE, we retained only LE44 as these are the basepairs that interact with the inner dimer. On the RE, as we did not know the optimal spacing between binding motifs, we generated a series of constructs that all included RE16 (i.e., the basepairs contacted by the core of the inner dimer) and the 19 bp sequence that comprises that pBac 2xCRD binding site, but in which the spacing between the two regions was increased from 0 to 9 bp by adding authentic RE sequence (generating RE16 through RE25). We reasoned that this modification would be well-tolerated given the dearth of protein-DNA contacts to the RE between the two dimers of the tetramer. As shown in Figure 7C, when cells were transfected with a plasmid encoding only one of the modified *pBat* transposases (“R” or “D” alone) and the pDonor plasmids, there was no detectable transposition activity. However, when both modified *pBat* transposase plasmids were co-transfected (“RD”), transposition activity was dependent on the spacing length with peak activity corresponding to RE21 and RE22. Under these experimental conditions, the robust heterodimer activity was ∼50% of that of hyperactive 4StoA pBat with LE88-RE100 transposon ends^12^ (green bar, Figure 7C). We conclude that this transposition activity was due to the “RD” heterodimers synapsing and integrating the redesigned TIRs.

### The pBat-2xCRD(D162R)/pBat-pB2xCRD(R160D) heterodimer can direct site-specific targeting

To test the ability of our redesigned heterodimer to serve as the foundation for developing site-directed integration systems, we took advantage of a recently described approach employing fusions of the *pBat* transposase with TALEs. Short et al.^22^ have demonstrated that a specific TTAA site in human chromosome 6 identified as a potential GSH can be targeted by pBat using one TALE designed to bind to a 9-bp sequence 18 bp upstream of the TTAA and a second TALE that binds 9-bp located 18 bp downstream. We incorporated their strategy into the heterodimer “R” and “D” proteins and retained their design principle of inserting the TALE domains between amino acids 64 and 65 of the disordered N-terminal domain; however, we assayed activity with a LE44-RE21pB2xCRD transposon rather than the full *pBat* transposon ends (Figure 7D).

Transposition targeting was initially assayed in a three-plasmid transposition experiment in which the 58-bp target human chr6 sequence was introduced into a pTarget plasmid (Figure 7E). pTarget contains five TTAA sites, and we assessed the ability of the modified heterodimers to direct integration from plasmid-to-plasmid into the chr6 site (designated “TTAA_6” in Figure 7E). As a control, we used an identical target plasmid in which the TALE binding site sequences were scrambled. When the two TALE binding site sequences were scrambled, integration occurred into all five TTAA sites in the target plasmid without particular preference (Figure 7F). However, when we used the TALE-inserted heterodimer that also contained the R336A Exc+Int-point mutant, 98.4% of integrations were directed to the chr6 site (Figure 7G).

We then asked if the TALE-modified heterodimer could direct integrations into chr6 of HEK293T cells. Transposition was assayed by transfecting cells with a donor LE44-RE21pB2xCRD transposon and helper plasmids expressing the TALE-heterodimer components, and selecting for puromycin-resistant colonies. After 15 days, colonies were collected, the combined genomic DNA isolated, and used as a template for gPCR with a chr6-specific primer and transposon-specific primers to detect genome-LE and genome-RE transposon junctions. The sequenced PCR products reveal that the LE44-RE21pB2xCRD transposon was accurately inserted into the intended target site in chr6 (Figure S11). We conclude that the heterodimers are active as fusion proteins and able to integrate into a desired genomic sequence, both in a target plasmid and in the human genome.

## Discussion

While transpososomes display remarkable architectural diversity,^23^ cut-and-paste DNA transpososomes always assemble as multimers composed of an even number of transposase protomers. There are examples for dimers (Tn*5*,^24^ Mos1^25^), tetramers (MuA,^26^ IS*21*^27^), and even octamers (Hermes^28^). These assemblies often display a low order - at least a pseudo - rotational symmetry, reflecting the fact that the LE and RE ends of the transposon are usually similar (and the TIRs are, by definition, approximate inverted repeats of each other). Furthermore, experimental structure determination has often relied on “symmetrized” TIRs to help obtaining a homogenous population in solution to facilitate the experiments, especially true for X-ray crystallographic experiments. Nevertheless, the active transpososome can be asymmetric due to the presence of additional transposon-encoded proteins in the assembly, as seen in the case of the TnsABCD transpososome.^29^ The RAG1/2 assembly that initiates V(D)J recombination, although no longer an active DNA transposase, is also asymmetric as it synapses conserved recombination signal sequences (RSSs) with 12 basepair spacer between them at one end and a 23 spacer at the other end, the so-called 12/23 rule.^30^ The three-dimensional structures of the RAG assembly^31,32^ reveal that the cellular protein HMGB1 is crucial to assure asymmetric binding. Even in these last two cases, the cores of the assemblies containing the transposase protomers have an approximate rotational symmetry.

The *pBat* transpososome structure breaks the paradigm as it has no rotational symmetry, reflecting the very different organization of binding motifs on the LE and RE. Furthermore, all tetrameric and higher order transpososomes with available experimental structures to date display protein/protein interactions between the constituent dimers of the assemblies and the *pBat* transpososome breaks this paradigm as well, as there are no protein/protein interactions between the dimers in the STC. The likely reason for the requirement for additional transposase protomers is to provide additional TIR binding strength and specificity.

Transpososomes with more than two transposase protomers face the same problem as pBat of an oversupply of active sites since all evidence suggests that only two are needed to catalyze the nuclease and transesterification steps. The key architectural problem then becomes to prevent the unused active sites from functioning, either through a structural change of the active site itself that prevents cleavage or strand transfer, or by keeping the active site away from DNA. In the tetrameric bacteriophage Mu transpososome,^26^ the distribution of the transposase binding motifs on the TIRs causes the unused active sites to be rotated away from the transposon DNA. This is no doubt aided by an assembly process which begins with transposase monomers, so that protomers can be wiggled into place as the tetramer assembles. pBat does not have this freedom as it is a dimer prior to TIR binding, and must rely on differences in DNA binding modes to keep the DNA backbone away from the unused active sites. The key is the large bend of the LE whereas the RE is essentially straight, thereby establishing different LE/RE relative configurations at the two dimers of the tetramer. This permits the inner dimer to deploy a binding mode different from the outer one to keep the phosphate backbone safely away from the outer dimer active sites.

We have previously shown that, although the *pBat* LE has three functional transposase dimer binding sites, optimal transposition activity in HEK293T cells is obtained when the innermost binding site is deleted; further deletion leads to a dramatic reduction in activity. The most parsimonious explanation of this necessity for the inner catalytically inactive dimer (i.e., the assembly of a tetramer) is its contribution to the synapse of the LE and RE, and in particular those of the two CRDs used to bind the second palindrome P2_LE_ on the LE (Figure 1E). To test this hypothesis, we generated two transposase variants (Figure 7A). In one, we linked two pBat CRDs to each other to allow binding of the P1_LE_ palindrome without the need for a second transposase protomer. We also generated a transposase where we replaced two linked pBat CRDs with two linked CRDs from pBac. When assayed with an appropriately modified RE that includes the pBac palindromic CRD binding site, on their own these proteins are inactive (Figure 7C). However, when cotransfected, transposition activity is recovered. These results are consistent with the assumption that in the context of the tetramer, the inner dimer in pBat is indeed needed only for synapse formation, and that by moving the CRD palindrome binding function to the outer dimer, an active heterodimeric transposase can be assembled.

The modifications that resulted in the obligatory heterodimeric transposase concept were guided by our near-atomic resolution structure of the active transpososome. The first was the identification of point mutations in the dimer interface that reduce the likelihood of homodimer formation. (Interestingly, our attempts to introduce similar mutations in pBac were largely unsuccessful as dimer interface mutations resulted in the loss of transposition activity. We note that there are differences between the dimer interfaces of pBac and pBat, despite the overall structural similarity and the relatively high sequence identity, with the buried ionic interactions placed differently. The second structure-guided modification effort was the identification of specific mutants that were defective in integration but with essentially wild-type excision activity. Finding Exc+Int-mutants in *piggyBac* superfamily members is more challenging than for other cut-and-paste transposases. The complication arises as residues involved in the superfamily-wide property of TTAA target recognition are the same as those involved in the hairpin intermediate TTAA recognition at excision. However, we have established that the R336A mutant has the desired Exc+Int-phenotype, likely because of reduced non-specific target binding activity. Again, the three-dimensional STC structure was essential in pinpointing R336 as an Exc+Int-candidate.

The successful conversion of the tetrameric transpososome into a obligatory heterodimeric one is a key step toward using *pBat* in applications that requires site-specific targeting. Conceptually, our approach is similar to the heterodimeric Fok1 restriction enzyme fused to site-specific DNA binding domains that can target specific DNA cleavage to a desired genomic location.^33,34^ A heterodimeric transposase allows the addition of two different target-sequence-binding domains directed to sequences flanking a chosen genomic TTAA site while removing complications inherent to a homo-tetramer which has unnecessary targeting domains appended to the inner dimer N-termini. As integration always occurs at TTAA, in principle targeting can be precise to nucleotide resolution. Furthermore, it is expected that heterodimers could direct integration in an orientation-specific way as target capture is no longer two-fold symmetric centered on the TTAA tetranucleotide. We have demonstrated that, in a plasmid-to-plasmid assay, two TALE domains can be used in conjunction with the *pBat* transposase to successfully direct targeting to a genomic sequence located in chr6 of the human genome (Figure 7G).

A number of different approaches to insertions of large DNA segments into the human genome are being developed. Undirected integration has proven useful for certain applications such as chimeric antigen receptor T-cell (CAR-T) therapies,^35^ although random integration by γ-retroviral and lentiviral vectors comes with risk.^36–39^ Transposon-based vectors also integrate randomly and can present similar dangers,^40^ so it is very clear that the safety profile of integration systems needs improvement. There are promising site-specific approaches including the use of site-specific recombinases such as large serine integrases in PASTE^41^ and PASSIGE,^42^ CRISPR-associated transposases (CAST),^43^ evoCAST,^44^ and RNA-directed transposons.^45^ With *pBac*, a very interesting approach has been developed that combines the activity of an Exc+Int-*pBac* transposase with fused, active Cas9 to achieve RNA-directed site-specific integration.^46^ While promising, all these approaches have drawbacks. For example, adapting bacterial systems to work in mammalian cells has challenges, and some of these approaches can create unwanted double-strand breaks. In principle, a heterodimeric mammalian transposase with two targeting domains engineered to integrate specifically - with nucleotide precision and with orientation specificity - could have substantial advantages for safe, site-directed integration in mammalian cells.

Finally, the heterodimeric approach solves a problem that limited the safety of targeted integration: the reliance on binding affinity over avidity. Systems that rely on a single targeting domain require high binding affinity to recognize the target sequence. However, high-affinity binders often possess sufficient non-specific affinity to trap the fusion protein on off-target sequences distributed across the human genome - acting as a “genomic sponge.” This trapping drives off-target integration events. In contrast, we envision tuning the binding affinity of the two targeting domains of our obligate heterodimer for moderate or low binding, preventing them from directing off-site integration events. High-affinity binding would then only be achieved through the increase in avidity that occurs when both domains recognize adjacent sites simultaneously. The final refinement of the system is that binding site recognition must be accompanied by a precisely placed TTAA between the binding sites. By structurally enforcing these two biological “AND” gates, the pBat heterodimer offers the prospect of a highly specific, directional integration platform for safe genomic integration.

### Limitations of the study

Here, we have solved the cryo-EM structure corresponding to only one state of the *piggyBat* transposition process: that after strand transfer has been accomplished. It is unknown if the observations and conclusions drawn here apply to earlier steps of transposon end pairing, cleavage, and hairpin opening and closing. We have only tested our obligatory heterodimer concept with one pair of TALEs and one chromosomal target site, so it will be of great interest to extend this work to other DNA-binding modules and other human safe harbor sites. Further work in other human cell types, including primary cells, is also warranted.

## RESOURCE AVAILABILITY

### Lead contact

Further information and requests for resources and reagents should be directed to and will be fulfilled by the lead contact, Fred Dyda.

### Materials availability

Plasmids generated in this study are readily available to the scientific community.

### Data and code availability

The customized Python code for plasmid analysis will be deposited and made publically available. For the piggyBat strand transfer complex structure, the accession numbers are EMDB-77018 and PDB 13ED. For the apo piggyBat (1-498) structure, the accession number is EMDB-77093.

## Acknowledgements

We are grateful to Jesse Owen for sharing information on TALE-transposase fusion design and results prior to publication. We thank Florencia Pratto for input into the development of the custom code for plasmid analysis, as well as Yanxiang Cui from the NIDDK cryo-EM facility, and Ulirch Baxa, Huaibin Wang, and Chih-Ta Chien from the MICEF cryo-EM facility for their assistance in cryo-EM data collection. We also thank Haotian Lei for productive discussions about cryo-EM data processing. This research was supported by the Intramural Research Program of the National Institute of Diabetes and Digestive and Kidney Diseases (NIDDK) within the National Institutes of Health (NIH). The contributions of the NIH authors are considered Works of the United States Government. The findings and conclusions presented in this paper are those of the authors and do not necessarily reflect the views of the NIH or the U.S. Department of Health and Human Services.

## Author information

Conceptualization, A.B.H., and F.D.; experimental studies, R.M., A.B.H., R.P.D., and A.P.; data analysis, R.M., A.B.H., R.P.D., A.P. and F.D.; supervision, A.B.H., and F.D.; manuscript writing, R.M., A.B.H., and F.D. with reviewing and editing input from all authors.

## Declaration of interests

A patent related to this work is in the process of being filed by the U.S. Government with F.D. and A.B.H. as inventors. They have no further financial interests to declare. The other authors declare no competing interests.

## Methods

### Plasmid constructs

Genes encoding the full-length pBat and pBat (1-498) transposases were codon-optimized for expression in EXPI293T cells and cloned into the pD2610 expression plasmid^12^ by Twist Biosciences (South San Francisco, CA). Transposase helper plasmids for the colony count and plasmid-to-plasmid assays were derived from pFV4a-piggyBat.^12^ Point mutant versions were either obtained from Genscript (Piscataway, NJ) or generated by the QuikChange method using primers obtained from Integrated DNA Technologies (IDT; Coralville, IA). The plasmid encoding 4StoA-pBat-pB2xCRD was synthesized by Genscript. Transposon donor plasmids for colony count assays contained a p2NGFPmini backbone,^12^ and were synthesized by Genscript, as was the donor plasmid pTet_LE44RE21pB used for the TALE plasmid-to-plasmid assays. pTarget plasmids used for plasmid-to-plasmid assay were generated by restriction cloning using gBlocks from IDT and a modified pHSG298 (Takara Bio) backbone. All plasmid sequences were confirmed by either Plasmidsaurus (Eugene, OR) or Quintara Biosciences (Frederick, MD) using Oxford Nanopore Technology. The sequences of all oligonucleotides used in this work are listed in Table 1.

**Table 1.**
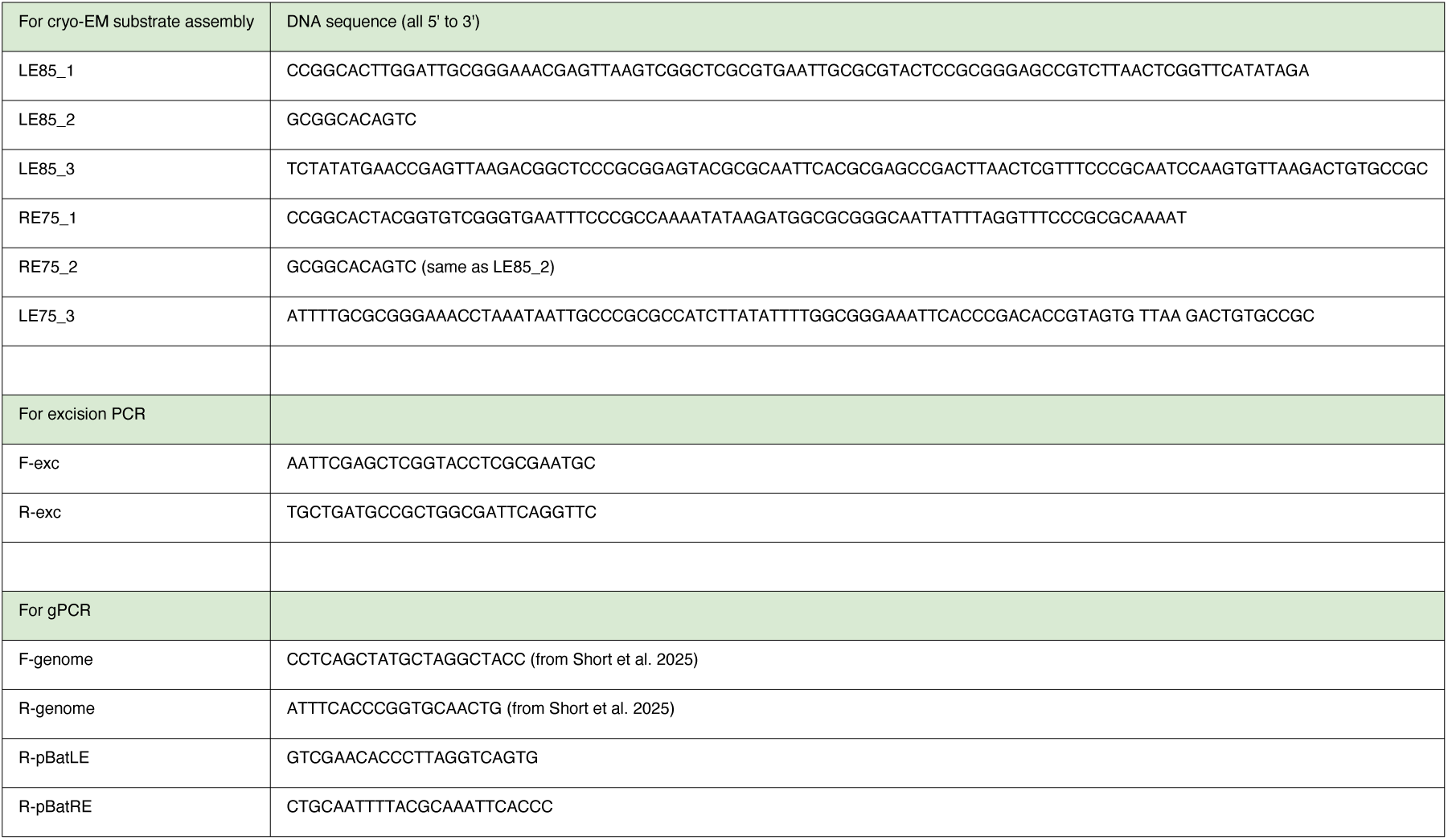
Oligonucleotides used in this work.

### Cell culture

HEK293T cells (ATCC #CRL-11268) were cultured under standard conditions using DMEM+Glutamax containing 10% (v/v) FBS and 0.1 mg/ml primocin (InvivoGen, San Diego, CA).

### Purification of full-length pBat and pBat (1-498)

Full-length pBat and pBat (1-498) were overexpressed and purified as described earlier^12^ with slight modifications. Briefly, plasmids pD2610-pBat and pD2610-pBat (1-498) were transfected into 500 ml of EXPI293F cells (ThermoFisher Scientific) using polyethylenimine (PEI), and cells were grown for 3 days at 37°C in a shaker incubator in the presence of 5% CO_2_. The cells were then harvested and stored at −80°C until use. For the purification of full-length pBat, the harvested cells were resuspended in a lysis buffer [50 mM Tris-HCl (pH 7.5), 500 mM NaCl, and 0.5 mM TCEP] containing DNase (Roche) and a protease inhibitor cocktail tablet (Roche). The cells were lysed by sonication and the cell lysates were centrifuged at 22,000 rpm for 45 min at 4°C. After centrifugation, the supernatant was collected, and mixed with 10 ml of amylose resin (New England BioLabs), equilibrated with the lysis buffer. The mixture was incubated for 1 h at 4°C on a rotating shaker and then loaded into a gravity flow column. The beads were washed with 100 ml of lysis buffer, and the fusion protein was eluted in buffer I [25 mM Tris-HCl (pH 7.5), 500 mM NaCl, 0.5 mM TCEP, and 10 mM maltose]. The eluted fusion protein was dialyzed overnight against dialysis buffer [50 mM Tris-HCl (pH 7.5), 500 mM NaCl, and 1 mM TCEP] at 4°C, and the MBP-tag was cleaved by TEV protease (250 U/mg recombinant protein) during dialysis. The sample was loaded onto a HiTrap Heparin HP column (Cytiva) and pBat transposase was eluted in buffer II [ 50 mM Tris-HCl (pH 7.5), 1 M NaCl, and 1 mM TCEP]. A final size-exclusion chromatography step was performed using a HiLoad Superdex 200 column (GE Healthcare, USA) run in buffer III [50 mM Tris-HCl (pH 7.5), 500 mM NaCl, and 1 mM TCEP].

A similar protocol was followed for the purification of pBat (1-498), with minor modifications in the purification steps after TEV protease cleavage. After TEV cleavage, the dialyzed sample was loaded onto a HiLoad Superdex 200 column run in buffer III. However, as MBP was not completely separated from the fractions containing pBat (1–498), a second affinity purification step was performed by mixing the pBat (1–498)-containing fractions with 1 ml of amylose resin equilibrated in lysis buffer, followed by incubation for 1 h at 4°C. The sample was then loaded onto a gravity flow column and purified using the same affinity purification protocol as above. pBat (1–498) was finally purified by size-exclusion chromatography on a HiLoad Superdex 200 column in buffer III.

### Preparation of strand transfer DNA

Oligonucleotides for the preparation of the strand transfer DNA were obtained from Integrated DNA Technologies (IDT; Coralville, IA) and resuspended in10 mM Tris-HCl pH 8.0 buffer. The LE85 strand transfer (LE85-ST) DNA was prepared by mixing an equimolar amount of (LE85_1) 5’-CCGGCAC TTGGATTGCGGGAAACGAGTTAAGTCGGCTCGCGTGAATTGCGCGTACTCCGCGGGAGCCGTCTTAACTCGGTTCATATAGA-3’; (LE85_2) 5’-GCGGCACAGTC-3’; and (LE85_3) 5’-TCTATATGAACCGAGTTAAGACGGCTCCCGCGGAGTACGCGCAATTCACGCGAGCCGA CTTAACTCGTTTCCCGCAATCCAAGTGTTAAGACTGTGCCGC-3’ at 80°C for 10 min, and then cooling overnight at room temperature to anneal. The same procedure was followed for assembling the RE75-ST DNA, where equimolar amounts of (RE75_1) 5’-CCGGCACTACGGTGTCGGGTGAATTTCCCGCCAAAATATAAGATGGCGCGGGCAATTATTTAGGTTTCCCGCGCAAAAT-3’; (RE75_2) 5’-GCGGCACAGTC-3’; and (RE75_3) 5’-ATTTTGCGCGGGAAACCTAAATAATTGCCCGCGCCATCTTATATTTTGGCGGGAAATTCACCCGACACCGTAGTG TTAA GACTGTGCCGC-3’ were mixed together and annealed as above.

### Assembly of pBat/LE85 ST/RE75 ST complex for Cryo-EM

The strand transfer complex was prepared by mixing 120 µM purified pBat with 24 µM LE85 ST DNA in buffer III. It was then incubated for 2 h at 4°C and dialyzed at 4°C against assembly buffer [25 mM HEPES (pH 7.5), 500 mM NaCl, 5 mM CaCl_2,_ and 1 mM TCEP] for 3 h. The complex was sequentially dialyzed against buffers with decreasing salt concentrations from 400 mM NaCl, 300 mM NaCl, 200 mM NaCl (3 h per step) and finally dialyzed in cryo-EM sample preparation buffer [25 mM HEPES (pH 7.5), 125 mM NaCl, 5 mM CaCl_2,_ and 1 mM TCEP]. After overnight dialysis, the complex was centrifuged for 10 min at 15,000 rpm at 4°C, and loaded onto a Superose 6 column (Cytiva), and was purified by size exclusion chromatography in the cryo-EM sample preparation buffer. To select fractions containing pBat bound to LE85 ST, eluted fractions were mixed with 50% glycerol (2.8 ml/10 ml sample) and loaded onto a 4% TBE-PAGE gel run for 1 h at 120 V at 4°C. The gel was stained with silver stain and imaged on an Azure 200 gel documentation system (Azure Biosystems). Both dimeric and tetrameric complexes were detected, and we selected fractions based on the abundance of the tetrameric species. The selected fractions were pooled together and concentrated using a 10 kDa cutoff Amicon Ultraconcentrator. The concentrate was mixed with 2 µl of 500 µM RE75 ST DNA and incubated overnight at 4°C. Finally, the assembled strand transfer complexes were applied to a Superose 6 column in cryo-EM sample preparation buffer, and the fractions were again analyzed using 4% TBE-PAGE.

### Cryo-EM sample preparation

Fractions containing pBat/LE85 ST/RE75 ST were collected and 2.9 µl of this sample at concentration ∼ 0.4 mg/ml (with A_280_:2.18 and A_260/280_:1.45) was applied immediately to a freshly glow discharged (30 sec at 15 mA, PELCO easiGlow) gold grid covered with a holey carbon film (Protochips C-Flat, 1.2/1.3, 300 mesh). After 5 sec wait time, it was blotted using the Vitrobot Mark IV (FEI) system for 4 sec (force 4) at 16°C and 100 % humidity, and was plunge-frozen in liquid ethane and transferred to liquid nitrogen for storage.

For pBat (1–498), an aliquot at 90 µM was loaded onto a Superose 6 column in cryo-EM sample buffer, and a fraction from the middle of the resulting peak was diluted with cryo-EM sample buffer to a final concentration of 0.25 mg/ml. For vitrification, we followed the same protocol as for the pBat/LE85 ST/RE75 ST complex with some minor modifications: 2.7 µl of sample at 0.25 mg/ml was applied on a freshly glow discharged gold grid covered with a holey carbon film (Protochips C-Flat, R1.2/1.3, 300 mesh) and plunge-frozen with the Vitrobot Mark IV (FEI) system in liquid ethane and stored in liquid nitrogen.

### Cryo-EM data collection

We have collected 8755 movies for the pBat/LE85 ST/RE75 ST complex using a Titan Krios G3 electron microscope (FEI, Thermo Fisher Scientific) at 300 kV, equipped with a K3 direct electron detector camera (Gatan) and an energy filter (20 eV slit width). The movies were recorded at 105,000 nominal magnification in super-resolution mode with a calibrated pixel size of 0.415 Å and a defocus range of −0.6 to −1.6 μm. The data was collected with the SerialEM program.^48^ Each movie had a total exposure time of 2.4 s with a total dose of 61.96 e^−^/Å^2^ per 40 frames.

The cryo-EM data of pBat (1–498) was collected similarly with some differences. The data was collected with a Titan Krios G4 electron microscope (FEI, Thermo Fisher Scientific), operating at 300 kV, and the microscope was equipped with an energy filter (20 eV slit width) and a K3 direct electron detector (Gatan). We collected a total of 14,416 movies, which were recorded at a nominal magnification of 105,000 in super-resolution mode with a calibrated pixel size of 0.412 Å, under a defocus range of −0.6 to −2.2 μm. The data was recorded with the EPU automation software (Thermo Fisher Scientific), and each movie consisted of 40 frames. The total exposure dose was 56.36 e^−^/Å^2^, with 1.8 s for each movie.

### Cryo-EM data analysis for the pBat/LE85 ST/RE75 ST complex

The cryo-EM images for the pBat/LE85 ST/RE75 ST complex were processed (see workflow in Figures S3 and S4) using the CryoSPARC v 4.6.0 software package.^49^ All the collected movies were imported, gain-corrected in CryoSPARC, followed by path motion correction and contrast transfer function (CTF) estimation. During the patch motion correction, the movies were binned by a factor of 2, resulting in a pixel size of 0.83 Å for subsequent data processing, and CTF parameters were estimated through patch-based CTF estimation. After CTF estimation, micrographs were denoised using the micrograph denoiser (using default parameters), and ∼ 1.4 million particles were picked from the randomly selected 3000 denoised micrographs using blob picker (100-300 Å particle diameter). The picked particles were manually filtered by adjusting the NCC score (>0.5) and power score (0-100). The particles (∼ 0.9 million) were extracted in a 480 x 480-pixel box size from the non-denoised micrographs and underwent 2D classification. While there was heterogeneity in this data set, it was dominated by tetrameric assemblies bound to two DNA oligonucleotides. There were also some pBat dimers with one DNA or two DNAs bound and also some DNA-unbound pBat dimers. Tetrameric assemblies of pBat bound to two DNAs were selected for further data processing. From the good 2D classes, 205,762 particles were subsequently used for ab-initio 3D reconstruction (four classes), and the best class (C2), containing 93,904 particles, was further selected for a round of 2D classification. After 2D classification, 53,910 particles were selected from the good 2D classes to generate templates, and particles were picked again from the same randomly selected 3000 denoised micrographs using the template picker. The picked ∼ 1.0 million particles were again manually filtered (NCC score: > 0.2 and power score: 20-150), and 843,864 particles extracted in a 500 x 500-pixel box size, which were used further for 2D classification.

The good 2D classes containing 414,516 particles were selected for heterogeneous refinement (default parameter), followed by ab-initio 3D reconstruction (three classes), and the 80,575 particles from the best class (C3) were selected again for a round of 2D classification. The 68,089 particles were selected from the best 2D classes to train Topaz 0.2.5a^50,51^ for the particle picking from 8,261 micrographs that eventually picked 877,779 particles, which were used for two rounds of heterogeneous refinement (default parameter with force hard classification) consecutively. From the second round of heterogeneous refinement, 308,472 particles were selected a further run of Topaz training and particle picking from all the micrographs. A total of ∼ 1.1 million particles were picked from all the micrographs, which were extracted in a 500 x 500-pixel box. The extracted particles were used for two rounds of heterogeneous refinement (default parameter with force hard classification), and after the second round, 325,891 particles were selected for ab-initio 3D reconstruction (two classes). The best class (C2) containing 187,903 particles was subjected to non-uniform refinement, which generated a map with a global resolution of 3.56 Å, as indicated by the gold-standard Fourier shell correlation (FSC) calculation at 0.143 FSC.

The particles from the non-uniform refinement were used for the reference-based motion correction in CryoSPARC v 4.6.0, followed again by non-uniform refinement, which produced a map with an overall resolution of 3.35 Å for the tetrameric assembly of pBat bound with two DNAs (Figures S4B-D). At this point, two overlapping masks were created separately for the individual pBat dimers bound with DNAs from the map obtained from non-uniform refinement (default parameter with threshold value, dilation radius, and soft padding width set to 0.013, 8 pixels, and 14 pixels, respectively). Focused refinements were performed for the two dimers separately, improving the resolutions to 2.86 Å and 2.93Å for the two dimers (Figures S4E-G, and S4H-J). Two focused maps were combined in Phenix v.2.0rc1-5555-000^52^ with the map obtained from the non-uniform refinement to generate a composite map, and this composite map was further sharpened and denoised with DeepEMhancer.^53^ This map was used for model building. The structure determination statistics are shown in Table 2.

**Table 2.** Cryo-EM data collection, refinement and validation statistics.

|  |  |
| --- | --- |
|  | piggyBat strand<br>transfer complex<br>(EMDB-77018)<br>(PDB 13ED) |
| **Data collection and processing** |  |
| Magnification | 105,000x |
| Voltage (kV) | 300 |
| Electron exposure (e-/Å<sup>2</sup>) | 61.96 |
| Defocus range (µm) | -0.6 to -1.6 |
| Pixel size (Å) | 0.415 Å |
| Symmetry imposed | none |
| Initial particle images (no.) | ~1,400,000 |
| Final particle images (no.) | 182,938 |
| Map resolution (Å) | 3.35 |
| FSC threshold | 0.143 |
| Map resolution range (Å) | 2.5-5 |
| **Refinement** |  |
| Initial model used (PDB code) | 9C0F |
| Model resolution (Å) | 2.5-5 |
| FSC threshold | 0.143 |
| Model resolution range (Å) | 2.5-5 |
| Map sharpening *B* factor (Å<sup>2</sup>) | -95.6 |
| Model composition |  |
| Non-hydrogen atoms | 22794 |
| Protein residues | 485+485+489+488 |
| DNA atoms | 1625+1861+163+163+1279+1469 |
| Ligands: Zn ion | 8 |
| Ca ion | 3 |
| *B* factors (Å<sup>2</sup>) |  |
| Protein | 109.43 |
| Nucleotides | 216.67 |
| Ligand | 197.25 |
| R.m.s. deviations |  |
| Bond lengths (Å) | 0.021 |
| Bond angles (°) | 1.812 |
| Validation |  |
| MolProbity score | 0.98 |
| Clashscore | 1.45 |
| Poor rotamers (%) | 0.11 |
| Ramachandran plot |  |
| Favored (%) | 97.51 |
| Allowed (%) | 2.43 |
| Disallowed (%) | 0.05 |

### Cryo-EM data processing for pBat (1**–**498)

The collected cryo-EM images of pBat (1–498) were processed (see workflow in Figures S9 and S10) with both CryoSPARC v 4.6.0 and RELION 5.0^54^ software packages. All 14,416 movies were imported and gain-corrected. The path motion correction and CTF estimation were performed in CryoSPARC v4.6.0 using the same protocol described earlier. In RELION 5.0, all the movies were motion-corrected with RELION’s own implementation, and the binning factor was set to 2, which provided a physical pixel of 0.824 Å for further processing. The CTF parameters were estimated using CTFFind 4.1.14^55^ in RELION 5.0.

In CryoSPARC, micrographs were manually curated, and those micrographs were discarded whose estimated CTF fit resolution was worse than 5 Å, relative ice thickness was over 1.2, and full frame motion distance was over 20 Å. We also removed other micrographs containing too much aggregation, excessive ice, or power spectrum artifacts. Further, micrographs were denoised using the same protocol described earlier, and approximately 1.7 million particles were picked from the randomly selected 1,000 denoised micrographs with blob picker (particle’s diameter = 50 - 100 Å). After particle picking, particles were inspected visually, and bad ones were removed by adjusting the NCC score > 0.8 and the power score = 5 - 15. Ultimately, ∼ 1.2 million particles were extracted from the micrographs in a 150 x 150-pixel box size and 2D classification performed (circular mask diameter = 120 Å and maximum alignment resolution = 6 Å).

After 2D classification, the particles from the ten good 2D classes were selected and used for ab-initio 3D reconstruction. The best class containing 202,003 particles was selected for a round of ab-initio 3D reconstruction (two classes). The particles from the best 3D class were extracted in a 300 x 300-pixel box size and used for non-uniform refinement. The map obtained from the non-uniform refinement was used to create fifty templates, and ∼ 4.6 million particles were picked from the randomly re-selected 3000 denoised micrographs with template picker. Particles were filtered by adjusting the NCC score > 0.5 and power score = 25-75. Approximately 3.2 million particles were extracted in a box size of 300 x 300 pixel and were used in heterogeneous refinement (default parameter with force hard classification).

Particles selected from the heterogeneous refinement were used in another round of 2D classification (small particle), and the parameters of circular mask diameter, maximum resolution, and maximum alignment resolution were set to 120 Å, 6 Å, and 6 Å, respectively. Good 2D classes were selected, and the particles from these classes were used for two rounds of ab-initio 3D reconstruction (four classes in each round) with an initial low-pass resolution of 12 Å and a maximum resolution of 4 Å. The three best classes (C2,C3,C4) were selected, and the particles from these classes were used for yet another round of 2D classification, using the same protocol described earlier. The good 2D classes containing 260,518 particles were selected for Topaz 0.2.5a for particle picking from all micrographs that picked ∼ 3.7 million particles, which were extracted in a 300 x 300-pixel box size. These particles were used for 2D classification (using the same parameter described earlier), and ∼ 3.6 million particles were selected from the good 2D classes, which were used further for three rounds of heterogeneous refinement (default parameter with force hard classification), followed by an ab-initio 3D reconstruction (three classes). The initial low-pass resolution and maximum resolution were set to 12 Å and 4 Å, respectively, for the ab-initio 3D reconstruction. The 3D map volume (C2) was selected from the ab-initio 3D reconstruction and used to create a solvent mask (default parameter with a dilatation radius of 6 pixels, soft padding width of 10 pixels, and threshold values of 0.07).

The selected particles from the ab-initio 3D reconstruction were subsequently used for 3D classification (three classes, with hard classification; filter resolution = 4 Å, initial structure lowpass resolution = 5 Å and initialization mode = PCA) and the best two classes (C2,C3) containing 273,071 particles were taken forward for non-uniform refinement, which provided a map with overall resolution of 2.91 Å resolution at 0.143 FSC, while applying C2 symmetry. CryoSPARC overestimated the map resolution, which was confirmed by manual inspection. Therefore, the particles from the non-uniform refinement were transferred into RELION 5.0 using PyEM^56^ for Bayesian polishing. After importing the particles into RELION 5.0, 273,071 particles were re-extracted from the motion corrected micrographs in RELION with a 300 x 300-pixel box size. The extracted particles were used for the 2D classification (mask diameter of 120 Å), and the best classes containing 219,403 particles were selected for 3D classification (three classes) using Blush regularization.^57^ After 3D classification, the best classes (C1 and C3) containing 149,073 particles were used and gold-standard-refined to 3.8 Å resolution at 0.143 FSC with blush regularization and C2 symmetry. After CTF refinement and particle polishing, the resolution was 3.3 Å, and RELION also overestimated the map resolution due to preferred orientation as the reconstruction did not allow the building of an atomic model.

### Atomic model building and refinement

DeepEMhancer was used to improve both the full and focused maps obtained after reference based motion correction and non uniform refinement. The two transposase dimers were manually docked using PDB code 9C0F^12^ while all the DNA was built fully manually using O Version 15.^58^ This was possible as the maps showed that essentially all basepairs were resolved so the DNA could be built into proper register. Where needed, the transposase was adjusted manually. Initially, the entire model was optimized with ChimeraX 1.8^59^ using real time molecular dynamics as implemented in Isolde^60^ and adjusted or rebuilt where needed. Further model refinement was carried out with Rosetta 2020.37.61417^61^ with the all-atom energy function set described in Alford et al.^62^ with increased Ramachandran weight (rama_prepro 0.7) using FastRelax in Cartesian space and b factor fitting. A composite map created by summing and normalizing (average 0.0, sd 1.0) the two focused maps in ChimeraX without any sharpening or other enhancements provided an additional energy term.^63^

### Excision assay

For a qualitative estimation of excision activity, 0.5 x 10^6^ HEK293T cells per well were used in a 6-well plate format. Approximately 18 hrs after initial plating, cells were transfected with 200 ng pTransposase (pFV4a-4StoA-pBat or mutants) and 400 ng pDonor (LE88-RE100) using Lipofectamine 3000 (Invitrogen/Thermo Fisher, Waltham, MA) according to the manufacturer’s instructions. Cells were harvested after 24 and 48 h, and low molecular weight plasmids recovered using a modified protocol with QIAprep Spin miniprep reagents (Qiagen, Hilden, Germany). Plasmids (5 μl) were used for PCR with 1X Q5 High-Fidelity DNA polymerase (NEB, Ipswitch, MA), 1X Q5 reaction buffer, 1X High GC Enhancer, 0.2 mM dNTP, and 0.4 μM each F-exc and R-exc primers as follows: 98°C 90 sec, then 27 cycles of 98°C 10 sec/60°C 20 sec/72°C 20 sec followed by 72°C for 120 sec. Samples were run on a 1% agarose/TAE gel containing SYBR Gold nucleic acid stain (Thermo Fisher). Visualization was with an Azure Biosystems 200 imager.

### Colony count integration assay

To measure integration activity of pBat point mutants, 0.5 x 10^6^ HEK293T cells per well were used in a 6-well plate format. Approximately 18 hrs after initial plating, cells were transfected with 100 ng pTransposase (pFV4a-4StoA-pBat or mutants) and 200 ng pDonor (LE88-RE100) using Lipofectamine 3000 according to the manufacturer’s instructions. The media was changed after 24 h. Cells were trypsinized after 48 h, and 1/20 of the cells re-plated on 10-cm plates in media containing 4 μg/ml puromycin. Media was changed every 3-4 days for 8-12 days. Cell were fixed with 4% formaldehyde in PBS, stained with 0.1% methylene blue, and manually counted.

To measure integration activity of the TALE constructs, the same procedure was followed except that cells were transfected with 20 ng each of the plasmids encoding the heterodimeric transposase and 10 ng pDonor.

### Plasmid-to-plasmid integration assay

The assay in HEK293T cells was performed similarly to the colony count assay except that cells were transfected with 0.5 μg each of the heterodimer plasmids, 1 μg of pDonor (LE44-RE21pB2xCRD), and 1.5 μg pTarget. Cells were harvested after 48 h, and low molecular weight plasmids recovered as described for the excision assay. The plasmid mix was digested with ScaI-HF (NEB) to linearize any unmodified pDonor plasmids, then purified using a QIAquick PCR purification kit (Qiagen) and 1 μl (after 1/3 dilution with water) was electroporated into 20 μl ElectroMAX DH10B cells (Invitrogen/Thermo Fisher) using a BioRAD Xcell Gene Pulser (pre-set E. coli program - 1 mm, 1.8 kV). Warm media (1 ml SOC) was added immediately, and cells grown 1 h at 37°C prior to plating the entire volume on 10-cm triple-resistance LB plates (30 μg/ml Kanamycin + 12 μg/ml Tetracycline + 30μg/ml Streptomycin; KDMedical, Columbia, MD) for 48 h at 30°C. Colonies were recovered by manually scraping all the cells from the plates, and low molecular weight plasmids isolated (QIAprep Spin miniprep kit). The plasmid mixes were then sequenced by commercial long read nanopore sequencing (Plasmidsaurus or QuintaraBio).

### Integration site analysis

We initially used the web-based OnRamp^64^ pipeline to assign individual nanopore reads from raw data fastq files to one of the 10 possible plasmid products of transposon integration into pTarget (five TTAA sites and two possible insertion orientations at each). We later used a custom Python script to assign reads based on a search for the unique transposon-plasmid transposon junctions corresponding to each of the ten possible plasmid products. Reads were assigned only (i) if they were within 250 bp of the correct size for pTarget containing a full-length transposon insertion; (ii) if both the LE and RE could be unambiguously detected; and (iii) the unique flanking sequences (including the TTAA TSD) could be assigned to a single TTAA site.

### gPCR to detect genomic insertions into human chr6

The transposition assay was carried out as described above, but after puro selection, cells were scraped from the plate and genomic DNA was isolated using a Monarch Spin gDNA Extraction kit (NEB). PCR reactions to detect genome-transposon junctions were carried out as follows: 1X Q5 High-Fidelity DNA polymerase, 1X Q5 reaction buffer, 0.2 mM dNTP, 100 ng gDNA, and 0.4 μM each F-genome and R-pBatLE or R-pBatRE primers as follows: 98°C 120 sec, then 40 cycles of 98°C 10 sec/67°C 20 sec/72°C 20 sec followed by 72°C for 120 sec. Products were separated and visualized on a 1.3% TAE gel, recovered using a QIAquick Gel Extraction Kit (Qiagen), and then ligated into pJET1.2 blunt (CloneJet, Thermo Scientific) according to the manufacturer’s instructions. Miniprep DNA from several of the resulting *E. coli* DH5α colonies were sequenced by long read nanopore and Sanger sequencing to confirm integration.

## Notes

### Competing Interest Statement

The authors have declared no competing interest.

## References

1. Liu, Y., Kong, J., Liu, G., Li, Z., and Xiao, Y. (2024). Precise gene knock-in tools with minimized risk of DSBs: A trend for gene manipulation. Adv. Sci. 11, 2401797. 10.1002/advs.202401797.

2. Hwang, J., Ye, D.Y., Jung, G.Y., and Jang, S. (2024). Mobile genetic element-based gene editing and genome engineering: Recent advances and applications. Biotechnol. Adv. 72, 108343. 10.1016/j.biotechadv.2024.108343.

3. Fong, J.H.C., and Ceroni, F. (2025). Transgene integration in mammalian cells: The tools, the challenges, and the future. Cell Syst. 16, 101426. 10.1016/j.cels.2025.101426.

4. Schmitz, M., and Querques, I. (2024). DNA on the move: mechanisms, functions and applications of transposable elements. FEBS Open Bio 14, 13–22. 10.1002/2211-5463.13743.

5. Chaaban, A., Sleem, R., Santina, J., Rima, M., and Ibrahim, J.-N. (2025). Exploring transposable elements: new horizons in cancer diagnostics and therapeutics. Mob. DNA 16, 28. 10.1186/s13100-025-00366-9.

6. Aznauryan, E., Yermanos, A., Kinzina, E., Devaux, A., Kapetanovic, E., Milanova, D., Church, G.M., and Reddy, S.T. (2022). Discovery and validation of human genomic safe harbor sites for gene and cell therapies. Cell Rep. Meth. 2, 100154. 10.1016/j.crmeth.2021.100154.

7. Wilson, M.H., Coates, C.J., and George, A.L. (2007). *PiggyBac* Transposon-mediated Gene Transfer in Human Cells. Mol. Ther. 15, 139–145. 10.1038/sj.mt.6300028.

8. Kovač, A., and Ivics, Z. (2017). Specifically integrating vectors for targeted gene delivery: progress and prospects. Cell Gene Ther. Insights 3, 103–123. DOI: 10.18609/cgti.2017.013.

9. Bhatt, S. and Chalmers, R. (2019). Targeted DNA transposition *in vitro* using a dCas9-transposase fusion protein. Nucleic Acids Res. 47, 8126–8135. 10.1093/nar/gkz552.

10. Ray, D.A., Feschotte, C., Pagan, H.J., Smith, J.D., Pritham, E.J., Arensburger, P., Atkinson, P.W., and Craig, N.L. (2008). Multiple waves of recent DNA transposon activity in the bat, *Myotis lucifugus*. Genome Res. 18, 717–728. 10.1101/gr.071886.107.

11. Mitra, R., Li, X., Kapusta, A., Mayhew, D., Mitra, R.D., Feschotte, C., and Craig, N.L. (2013). Functional characterization of *piggyBat* from the bat *Myotis lucifugus* unveils an active mammalian DNA transposon. Proc. Natl. Acad. Sci. U.S.A. 110, 234–239. 10.1073/pnas.1217548110.

12. Hickman, A.B., Lannes, L., Furman, C.M., Hong, C., Franklin, L., Ghirlando, R., Ghosh, A., Luo, W., Konstantinidou, P., Lorenzi, H.A., et al. (2025). Activity of the mammalian DNA transposon *piggyBat* from *Myotis lucifugus* is restricted by its own transposon ends. Nat. Commun. 16, 458. 10.1038/s41467-024-55784-9.

13. Chen, Q., Luo, W., Veach, R.A., Hickman, A.B., Wilson, M.H., and Dyda, F. (2020). Structural basis of seamless excision and specific targeting by *piggyBac* transposase. Nat. Commun. 11, 3446. 10.1038/s41467-020-17128-1.

14. Ivančić, D., Agudelo, A., Lindstrom-Vautrin, J., Jaraba-Wallace, J., Gallo, M., Das, R., Ragel, A., Herrero-Vicente, J., Higueras, I., Billeci, F., et al. (2025). Discovery and protein language model-guided design of hyperactive transposases. Nat. Biotechnol. 10.1038/s41587-025-02816-4.

15. Li, X., Burnight, E.R., Cooney, A.L., Malani, N., Brady, T., Sander, J.D., Staber, J., Wheelan, S.J., Joung, J.K., McDray, Jr., P.B., et al. (2013). *piggyBac* transposase tools for genome engineering. Proc. Natl. Acad. Sci. USA 110, E2279–E2287. 10.1073/pnas.1305987110.

16. Kesselring, L., Miskey, C., Zuliani, C., Querques, I., Kapitonov, V., Laukó, A., Fehér, A., Palazzo, A., Diem, T., Lustig, J., et al. (2020). A single amino acid switch converts the *Sleeping Beauty* transposase into an efficient unidirectional excisionase with utility in stem cell reprogramming. Nucleic Acids Res. 48, 316–331. 10.1093/nar/gkz1119.

17. Luo, W., Hickman, A.B., Genzor, P., Ghirlando, R., Furman, C.M., Menshikh, A., Haase, A., Dyda, F., and Wilson, M.H. (2022). Transposase N-terminal phosphorylation and asymmetric transposon ends inhibit *piggyBac* transposition in mammalian cells. Nucleic Acids Res. 50, 13128–13142. 10.1093/nar/gkac1191.

18. Baker, N.A., Sept, D., Joseph, S., Holst, M.J., and McCammon, J.A. (2001). Electrostatics of nanosystems: application to microtubules and the ribosome. Proc. Natl. Acad. Sci. U.S.A. 98, 10037–10041. 10.1073/pnas.181342398.

19. Wang, K., Hu, G., Basu, S., and Kurgan, L. (2024). flDPnn2: Accurate and Fast Predictor of Intrinsic Disorder in Proteins. J. Mol. Biol. 436, 168605. 10.1016/j.jmb.2024.168605.

20. Duvaud, S., Gabella, C., Lisacek, F., Stockinger, H., Ioannidis, V., and Durinx, C. 2021. Expasy, the Swiss Bioinformatics Resource Protal, as designed by its users. Nucleic Acids Res. 49, W216–W227. 10.1093/nar/gkab225.

21. Olsson, M.H.M., Søndergaard, C.R., Rostkowski, M., and Jensen, J.H. (2011). PROPKA3: Consistent treatment of internal and surface residues in empirical pKa predictions. J. Chem. Theory Comput. 7, 525–537. 10.1021/ct100578z.

22. Short, J.E., Sharek, L., Meckler, J.F., Stoytchev, I., Tran, C.T., Errard, C., Hew, B.E., Johnson, B.E., Waller, D.F., Xie, S., et al. (2025). Gene-sized DNA insertion at genomic safe harbors in human cells using a site-directed transposase. Nucleic Acids Res. 53, gkaf1316. 10.1093/nar/gkaf1316.

23. Dyda, F., Chandler, M., and Hickman, A.B. (2012). The emerging diversity of transpososome architectures. Quart. Rev. Biophys. 45, 493–521. 10.1017/S0033583512000145.

24. Davies, D.R., Goryshin, I.Y., Reznikoff, W.S., and Rayment, I. (2000). Three-dimensional structure of the Tn*5* synaptic complex transposition intermediate. Science 289, 77–85. 10.1126/science.289.5476.77.

25. Richardson, J.M., Colloms, S.D., Finnegan, D.J., and Walkinshaw, M.D. (2009). Molecular Architecture of the Mos1 Paired-End Complex: The Structural Basis of DNA Transposition in a Eukaryote. Cell 138, 1096–1108. 10.1016/j.cell.2009.07.012.

26. Montaño, S., Pigli, Y., and Rice, P. (2012). The Mu transpososome structure sheds light on DDE recombinase evolution. Nature 491, 413–417. 10.1038/nature11602.

27. de la Gándara, Á., Spínola-Amilibia, M., Araújo-Bazán, L., Núñez-Ramírez, R., Berger, J.M., and Arias-Palomo, E. (2024). Molecular basis for transposase activation by a dedicated AAA+ ATPase. Nature 630, 1003–1011. 10.1038/s41586-024-07550-6.

28. Lannes, L., Furman, C.M., Hickman, A.B., and Dyda, F. (2023). Zinc-finger BED domains drive the formation of the active *Hermes* transpososome by asymmetric DNA binding. Nat. Commun. 14, 4470. 10.1038/s41467-023-40210-3.

29. Wang, S., Siddique, R., Hall, M.C., Rice, P.A., and Chang, L. (2024). Structure of TnsABCD transpososome reveals mechanisms of targeted DNA transposition. Cell 187, 6865–6881.e16. 10.1016/j.cell.2024.09.023.

30. Tonegawa, S. (1983). Somatic generation of antibody diversity. Nature 302, 575–581. 10.1038/302575a0.

31. Ru, H., Chambers, M.G., Fu, T-M., Tong, A.B., Liao, M., and Wu, H. (2015). Molecular Mechanism of V(D)J Recombination from Synaptic RAG1-RAG2 Complex Structures. Cell 163, 1138–1152. 10.1016/j.cell.2015.10.055.

32. Chen, X., Cui, Y., Wang, H., Zhou, Z.H., Gellert, M., and Yang, W. (2020). How mouse RAG recombinase avoids DNA transposition. Nat. Struct. Mol. Biol. 27, 127–133. 10.1038/s41594-019-0366-z.

33. Miller, J.C., Holmes, M.C., Wang, J., Guschin, D.Y., Lee, Y.L., Rupniewski, I., Beausejour, C., Waite, A.J., Wang, N.S., Kim, K.A., et al. (2007). An improved zinc-finger nuclease architecture for highly specific genome editing. Nature Biotechnol. 25, 778–785. 10.1038/nbt1319.

34. Szczepek, M., Brondani, V., Büchel, J., Serrano, L., Segal, D.J., and Cathomen, T. (2007). Structure-based redesign of the dimerization interface reduces the toxicity of zinc-finger nucleases. Nature Biotechnol. 25, 786–793. 10.1038/nbt1317.

35. Labanieh, L., and Mackall, C.L. (2023). CAR immune cells: design principles, resistance and the next generation. Nature 614, 635–648. 10.1038/s41586-023-05707-3.

36. https://www.fda.gov/vaccines-blood-biologics/safety-availability-biologics/fda-requires-boxed-warning-t-cell-malignancies-following-treatment-bcma-directed-or-cd19-directed

37. Liao, Q., and Xu, J. (2025). Safe CAR-T: shedding light on CAR-related T-cell malignancies. EMBO Mol. Med. 17, 589–593. 10.1038/s44321-025-00205-7.

38. Deichmann, A., Hacein-Bey-Abina, S., Schmidt, M., Garrigue, A., Brugman, M.H., Hu, J., Glimm, H., Gyapay, G., Prum, B., Fraser, C.C., et al. (2007). Vector integration is nonrandom and clustered and influences the fate of lymphopoiesis in SCID-X1 gene therapy. J. Clin. Invest. 117, 2225–2232. 10.1172/JCI31659.

39. Fraietta, J.A., Lacey, S.F., Orlando, E.J., Pruteanu-Malinici, I., Gohil, M., Lundh, S., Boesteanu, A.C., Wang, Y., O’Connor, R.S., Hwang, W.T., et al. (2018). Determinants of response and resistance to CD19 chimeric antigen receptor (CAR) T cell therapy of chronic lymphocytic leukemia. Nat. Med. 24, 563–571. 10.1038/s41591-018-0010-1.

40. Micklethwaite, K.P., Gowrishankar, K., Gloss, B.S., Li, Z., Street, J.A., Moezzi, L., Mach, M.A., Sutrave, G., Clancy, L.E., Bishop, D.C., et al. (2021). Investigation of product-derived lymphoma following infusion of *piggyBac*-modified CD19 chimeric antigen receptor T cells. Blood 138, 1391–1405. 10.1182/blood.2021010858.

41. Yarnall, M.T.N., Ioannidi, E.I., Schmitt-Ulms, C., Krajeski, R.N., Lim, J., Villiger, L., Zhou, W., Jiang, K., Garushyants, S.K., Roberts, N., et al. (2023). Drag-and-drop genome insertion of large sequences without double-strand DNA cleavage using CRISPR-directed integrases. Nat. Biotechnol. 41, 500–512. 10.1038/s41587-022-01527-4.

42. Pandey, S., Gao, X.D., Krasnow, N.A., McElroy, A., Tao, Y.A., Duby, J.E., Steinbeck, B.J., McCreary, J., Pierce, S.E., Tolar, J., et al. (2025). Efficient site-specific integration of large genes in mammalian cells via continuously evolved recombinases and prime editing. Nat. Biomed. Eng. 9, 22–39. 10.1038/s41551-024-01227-1.

43. Klompe, S.E., Vo, P.L.H., Halpin-Healy, T.S., and Sternberg, S.H. (2019). Transposon-encoded CRISPR-Cas systems direct RNA-guided DNA integration. Nature 571, 219–225. 10.1038/s41586-019-1323-z.

44. Witte, I.P., Lampe, G.D., Eitzinger, S., Miller, S.M., Berríos, K.N., McElroy, A.N., King, R.T., Stringham, O.G., Gelsinger, D.R., Vo, P.L.H., et al. (2025). Programmable gene insertion in human cells with a laboratory-evolved CRISPR-associated transposase. Science 388, eadt5199. 10.1126/science.adt5199.

45. Perry, N.T., Bartie, L.J., Katrekar, D., Gonzalez, G.A., Durrant, M.G., Pai, J.J., Fanton, A., Martins, J.Q., Hiraizumi, M., Ricci-Tam, C., et al. (2026). Megabase-scale human genome rearrangement with programmable bridge recombinases. Science 391, adz0276. 10.1126/science.adz0276.

46. Pallarès-Masmitjà, M., Ivančić, D., Mir-Pedrol, J., Jaraba-Wallace, J., Tagliani, T., Oliva, B., Rahmeh, A., Sánchez-Mejías, A., and Güell, M. (2021). Find and cut-and-transfer (FiCAT) mammalian genome engineering. Nat. Commun. 12, 7071. 10.1038/s41467-021-27183-x.

47. Dolinsky, T.J., Nielsen, J.E., McCammon, J.A., and Baker, N.A. (2004) PDB2PQR: an automated pipeline for the setup, execution, and analysis of Poisson-Boltzmann electrostatics calculations. Nucleic Acids Res. 32, W665–W667. 10.1093/nar/gkh381.

## Methods references

48. Mastronarde, D.N. (2005). Automated electron microscope tomography using robust prediction of specimen movements. J. Struct. Biol. 152, 36–51.

49. Punjani, A., Rubinstein, J.L., Fleet, D.J., and Brubaker, M.A. (2017). cryoSPARC: algorithms for rapid unsupervised cryo-EM structure determination. Nat. Meth. 14, 290–296. 10.1038/nmeth.4169.

50. Bepler, T., Morin, A., Rapp, M., Brasch, J., Shapiro, L., Noble, A.J., and Berger, B. (2019). Positive-unlabeled convolutional neural networks for particle picking in cryo-electron micrographs. Nat. Meth. 16, 1153–1160. 10.1038/s41592-019-0575-8.

51. Bepler, T., Kelley, K., Noble, A.J., and Berger, B. (2020). Topaz-Denoise: general deep denoising models for cryoEM and cryoET. Nat. Comm. 11, 5208. 10.1038/s41467-020-18952-1.

52. Liebschner, D., Afonine, P.V., Baker, M.L., Bunkóczi, G., Chen, V.B., Croll, T.I., Hintze, B., Hung L.W., Jain, S., McCoy, A.J., et al. (2019). Macromolecular structure determination using X-rays, neutrons and electrons: recent developments in Phenix. Acta Crystallogr D 75, 861–877. 10.1107/S2059798319011471.

53. Sanchez-Garcia, R., Gomez-Blanco, J., Cuervo, A., Carazo, J.M., Sorzano, C.O.S., and Vargas, J. (2021). DeepEMhancer: a deep learning solution for cryo-EM volume post-processing. Commun. Biol. 4, 874. 10.1038/s42003-021-02399-1.

54. Burt, A., Toader, B., Warshamanage, R., von Kügelgen, A., Pyle, E., Zivanov, J., Kimanius, D., Bharat, T.A.M., and Scheres, S.H.W. (2024). An image processing pipeline for electron cryo-tomography in RELION-5. FEBS Open Bio. 14, 788–1804. 10.1002/2211-5463.13873.

55. Rohou, A., and Grigorieff, N. (2015). CTFFIND4: Fast and accurate defocus estimation from electron micrographs. J. Struct. Biol. 192, 216–221. 10.1016/j.jsb.2015.08.008.

56. Kim, J., and Kang, J.Y. (2024). Practices in cryo-EM single particle analysis data processing: data transfer between RELION and cryoSPARC. Biodesign 12, 29–38. 10.34184/kssb.2024.12.2.29.

57. Kimanius, D., Jamali, K., Wilkinson, M.E., Lövestam, S., Velazhahan, V., Nakane, T., and Scheres, S.H.W. (2024). Data-driven regularization lowers the size barrier of cryo-EM structure determination. Nat. Meth. 21, 1216–1221. 10.1038/s41592-024-02304-8.

58. Jones, T.A., and Kjeldgaard, M. (1997). Electron-density map interpretation. Meth. Enzymol. 277, 173–208. 10.1016/S0076-6879(97)77012-5.

59. Meng, E.C., Goddard, T.D., Pettersen, E.F., Couch, G.S., Pearson, Z.J., Morris, J.H., and Ferrin, T.E. (2023). UCSF ChimeraX: Tools for structure building and analysis. Protein Sci. 32, e4792. 10.1002/pro.4792.

60. Croll, T.I. (2018). ISOLDE: a physically realistic environment for model building into low-resolution electron-density maps. Acta Crystallogr D 74, 519–530. 10.1107/S2059798318002425.

61. Leman, J.K., Weitzner, B.D., Lewis, S.M., Adolf-Bryfogle, J., Alam, N., Alford, R.F., Aprahamian, M., Baker, D., Barlow, K.A., Barth, P., et al. (2020). Macromolecular modeling and design in Rosetta: recent methods and frameworks. Nat. Meth. 17, 665–680. 10.1038/s41592-020-0848-2.

62. Alford, R.F., Leaver-Fay, A., Jeliazkov, J.R., O’Meara, M.J., DiMaio, F.P., Park, H., Shapovalov, M.V., Renfrew, P.D., Mulligan, V.K., Kappel, K., et al. (2017). The Rosetta All-Atom Energy Function for Macromolecular Modeling and Design. J. Chem. Theory Comput. 13, 3031–3048. 10.1021/acs.jctc.7b00125.

63. Wang, R.Y., Song, Y., Barad, B.A., Cheng, Y., Fraser, J.S., DiMaio, F., and Brunger, A.T. (2016). Automated structure refinement of macromolecular assemblies from cryo-EM maps using Rosetta. eLife 5, 17219. 10.7554/eLife.17219.

64. Mumm, C., Drexel, M.L., McDonald, T.L., Diehl, A.G., Switzenberg, J.A., and Boyle, A.P. 2023. Multiplexed long-read plasmid validation and analysis using OnRamp. Gen. Res. 33, 741–749. 10.1101/gr.277369.122.

