## Supplemental Figures for "Structure-based recasting of a mammalian DNA transpososome as an obligate heterodimer"

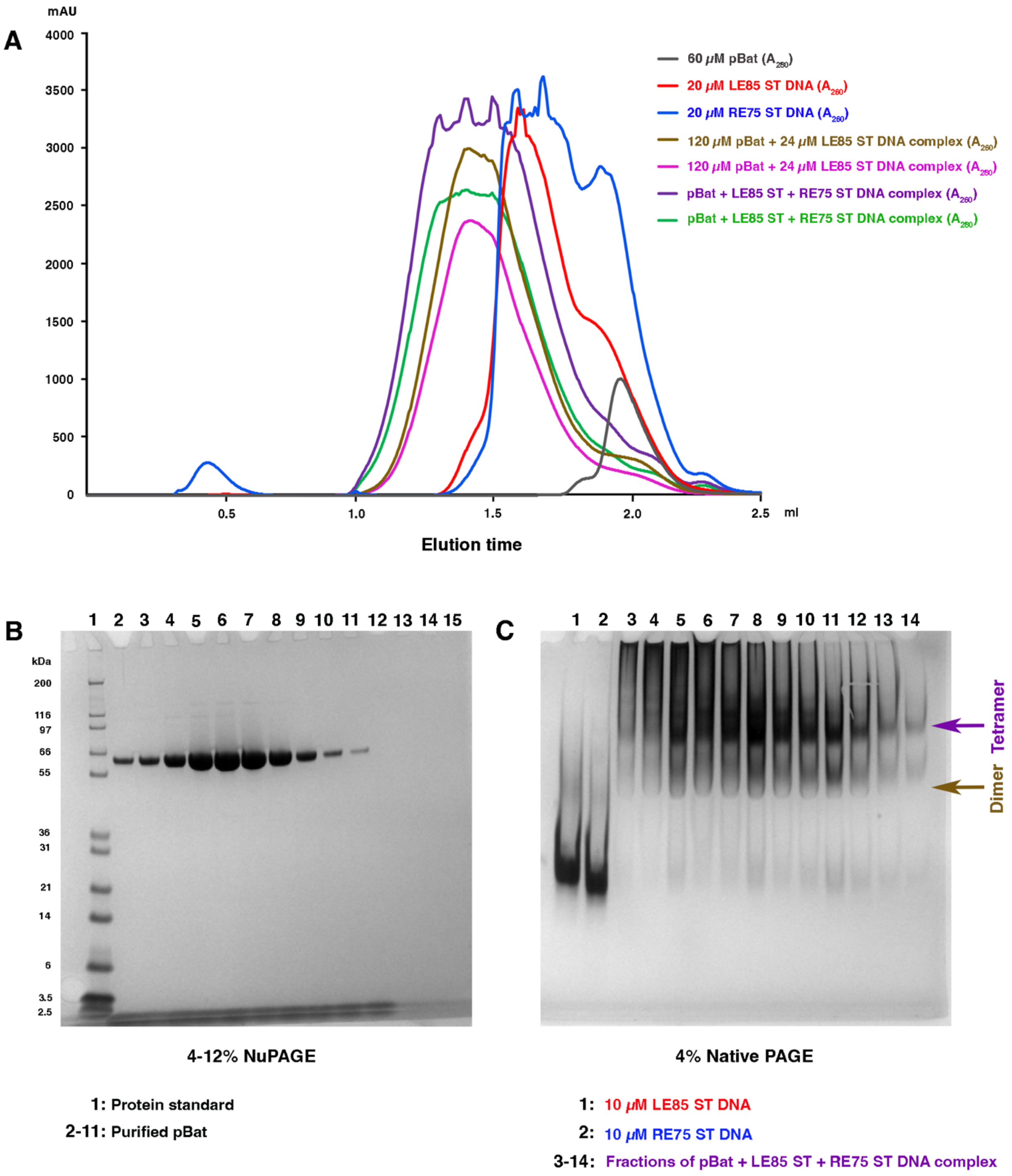


**Supplementary Figure 1. Preparation of pBat STC assembly.** (A) The chromatogram represents the elution profiles of pBat ST complex [pBat in complex with both LE 85 ST and RE75 ST DNA; purple (260 nm); green (280 nm)], along with the pBat:LE85 ST DNA complex [tan (260 nm); pink (280 nm)], as well as pBat [gray (280 nm)], LE85 ST DNA [red (260 nm)] and RE75 ST DNA [blue (260 nm)]. (B) NuPAGE analysis of purified pBat used for the cryo-EM sample preparation. (C) Native PAGE analysis of the purified pBat STC on a 4% polyacrylamide gel. Although both dimer and tetrameric species were present in the sample, the tetrameric species was vastly predominant.

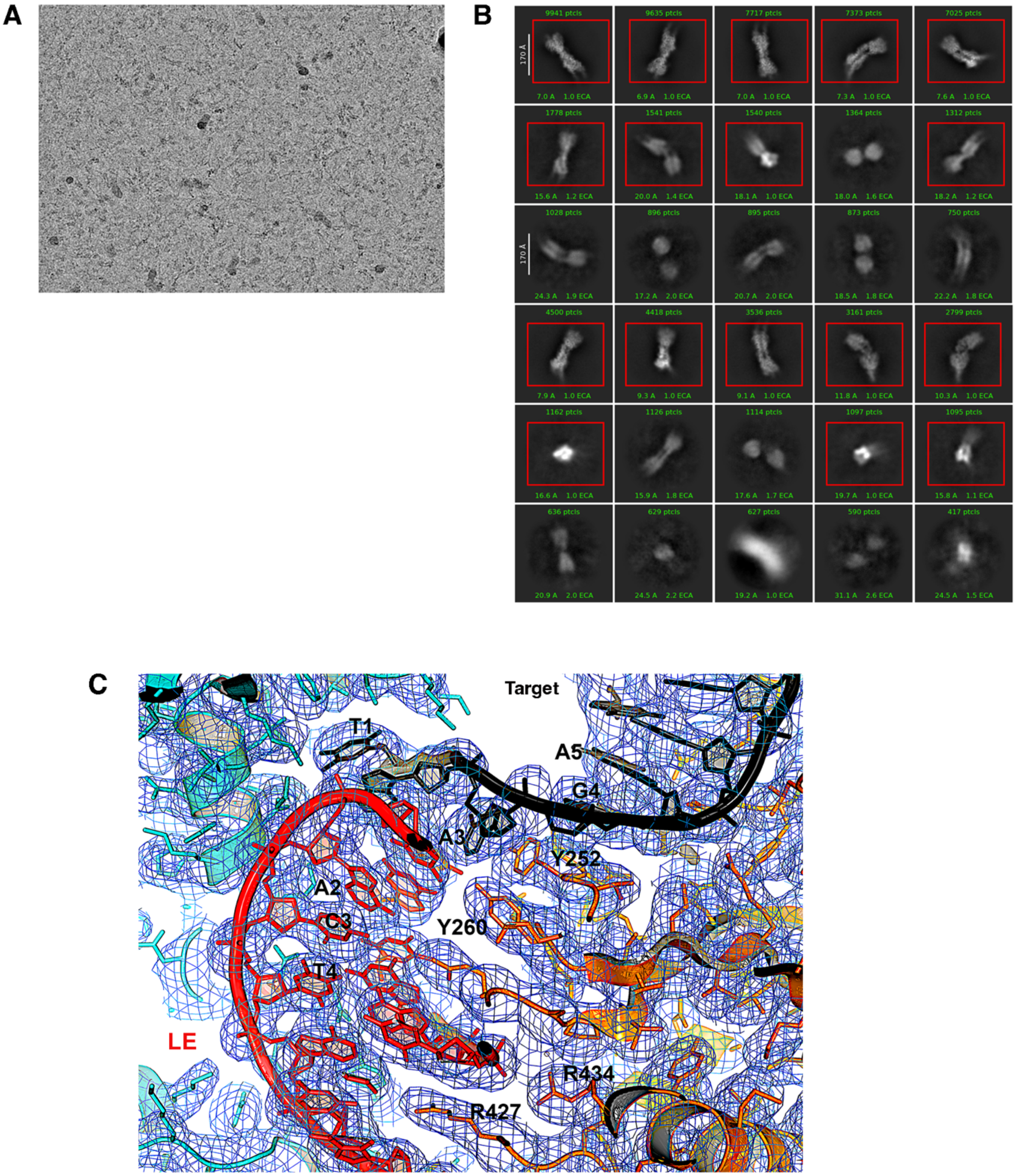


**Supplementary Figure 2. Sample micrograph, 2D class averages, and consistency of the built model with the electrostatic potential density map.** (A) Example of a motion-corrected micrograph of the pBat STC assembly. (B) The 2D class averages of the particles from pBat STC, showing a dominant crescent-shaped projection of tetrameric species. The red colored boxes indicate that particles from these classes were selected for cryo-EM data processing. (C) A view of the electrostatic potential density after DeepEMhancer. The map was normalized and contoured at the 3 sigma level.

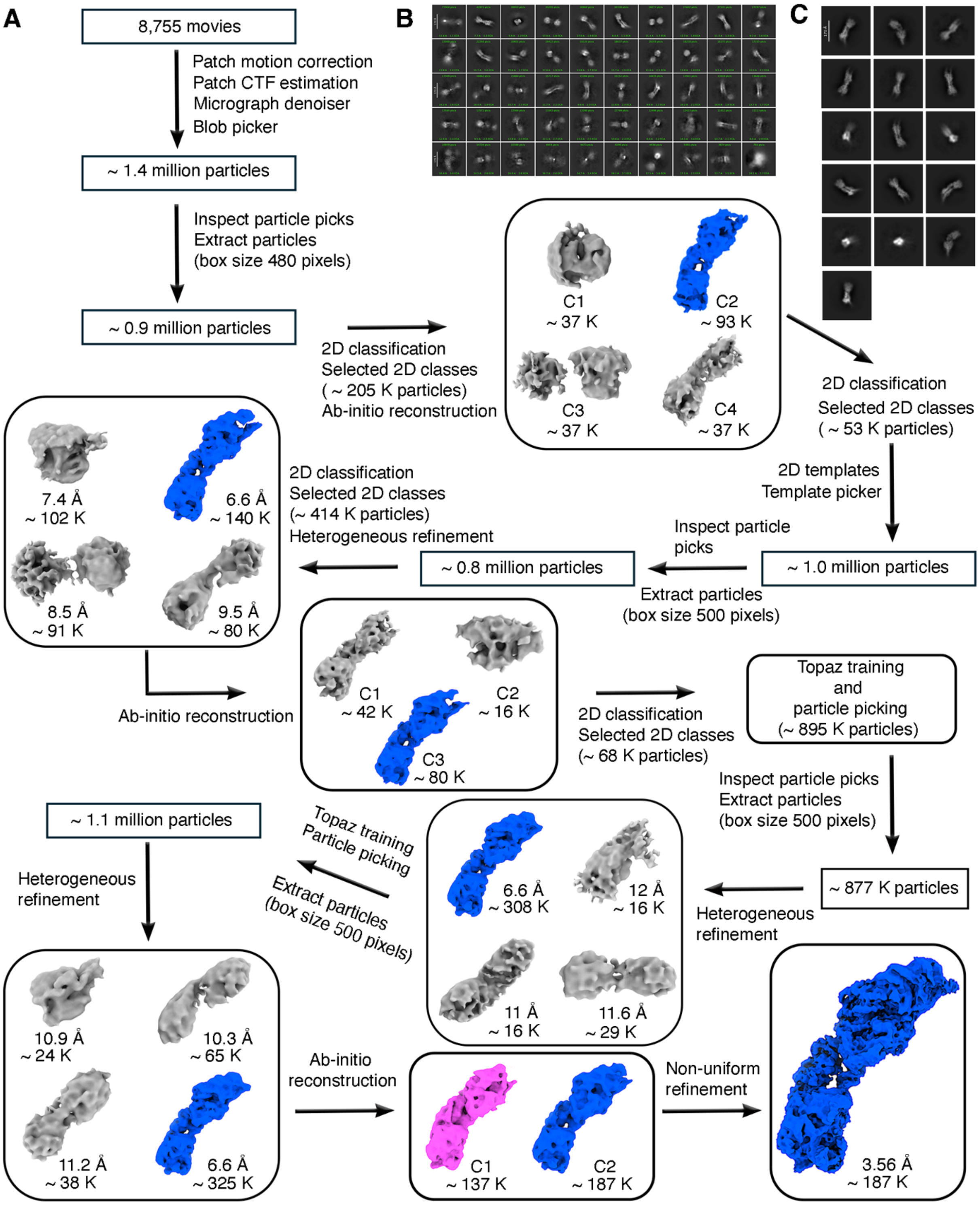


**Supplementary Figure 3. Cryo-EM data processing workflow of pBat STC**. (A) Detailed cryo-EM data processing workflow. Only the particles associated with the blue color map were selected for the next step of data processing from the earlier step, and the remaining particles were not considered for data processing. (B) Reference-free 2D class averages of the pBat STC in the early stage of data processing. While there was some heterogeneity present in the data set, it was predominated by the crescent-shaped tetrameric species bound to two DNA molecules. (C) The selected 2D class averages in the later stage of data processing.

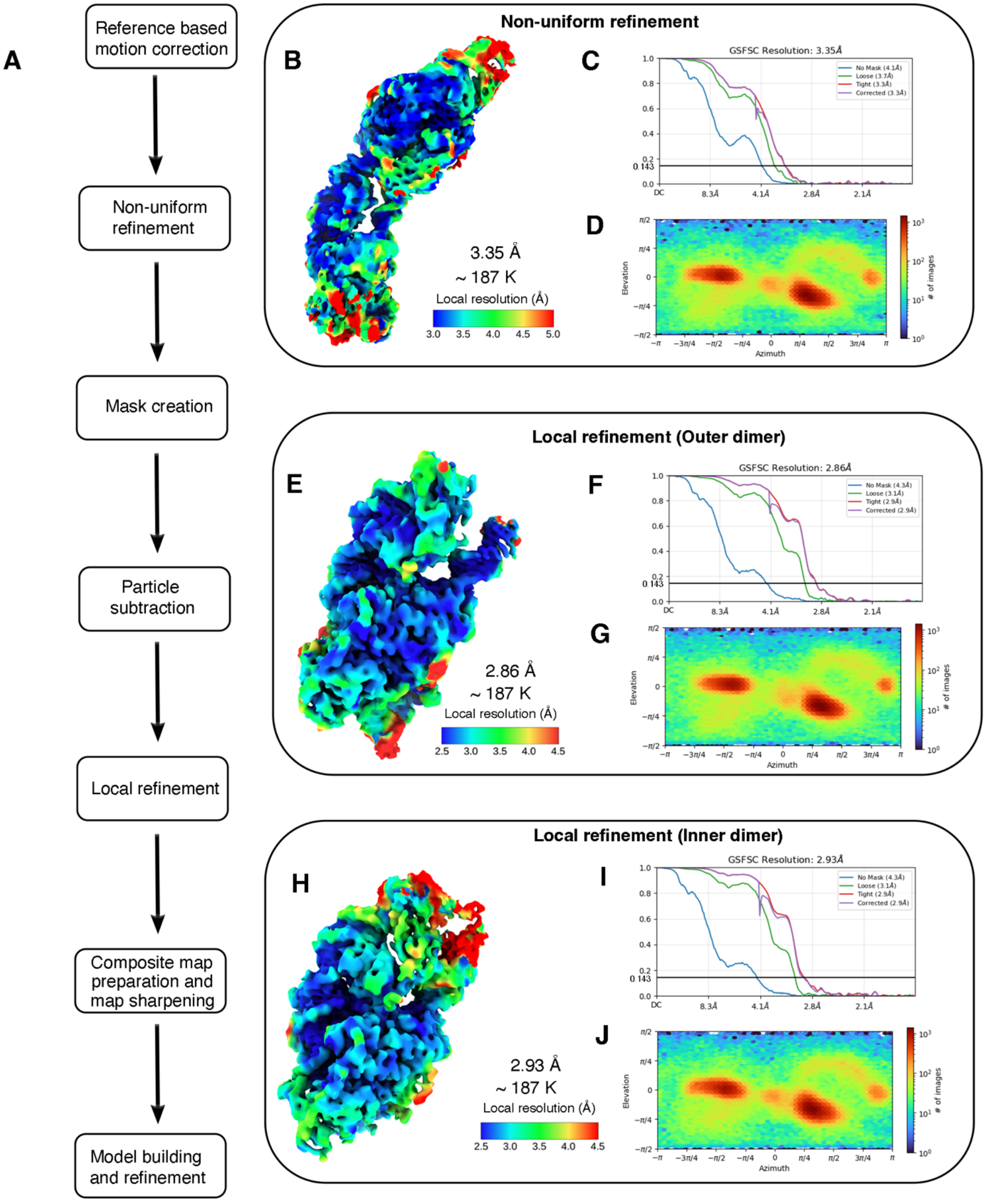


**Supplementary Figure 4. Cryo-EM data processing workflow of pBat STC**. (A) Detailed post-processing workflow in CryoSPARC. (B) Resolution distribution of the final cryo-EM map obtained from the non-uniform refinement in CryoSPARC. The contour was set to 0.03. (C) The final map achieved a resolution of 3.35 Å at 0.143 FSC. (D) Directional distribution of the particles viewed. (E) Resolution distribution of the focused map (outer dimer) obtained from local refinement in CryoSPARC. The contour was set to 0.03. (F) The final resolution of the focused map was achieved at 2.86 Å at 0.143 FSC. (G) Directional distribution of the particles viewed. (H) Resolution distribution of the focused map (inner dimer) obtained from local refinement in CryoSPARC at a contour of 0.03. (I) The focused map resolution was achieved at 2.93 Å at a 0.143 FSC for the inner dimer. (J) Directional distribution of the particles viewed.

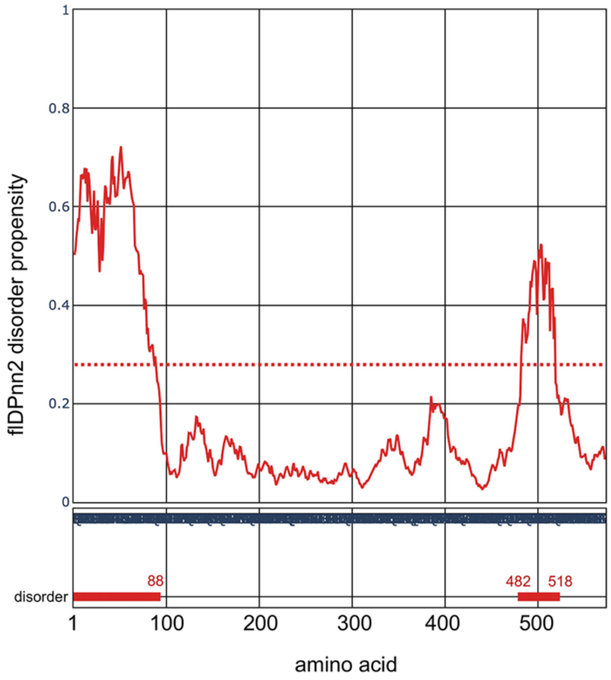


**Supplementary Figure 5.** Result of the "putative function- and linker-based Disorder Prediction using deep neural network 2” (flDPnn2) server used to predict intrinsically disordered regions in proteins. The output indicates that amino acids 1-88 and 482-518 of the pBat transposase are predicted to be disordered.

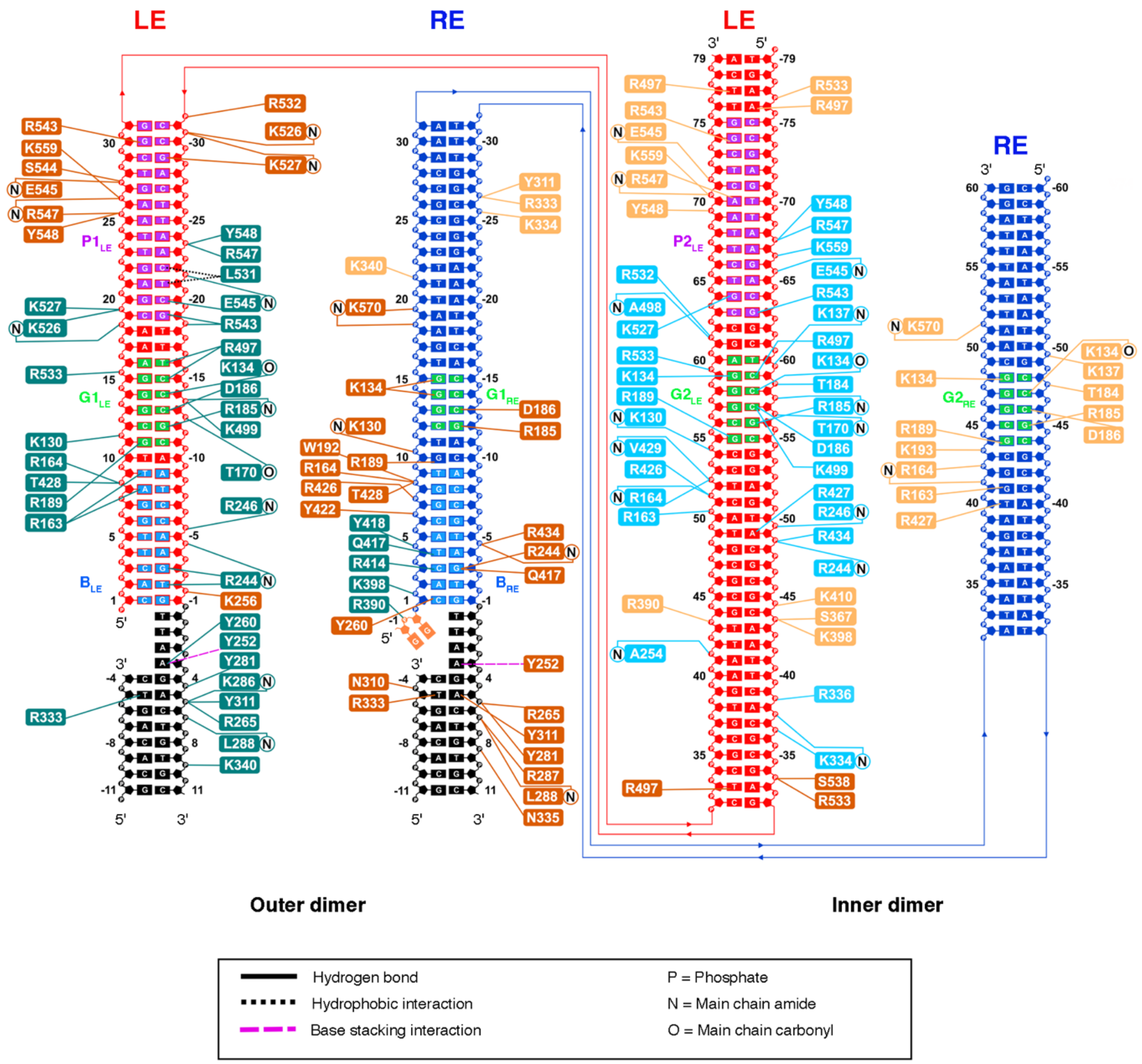


**Supplementary Figure 6. Schematic of interactions between pBat and DNA.** The colors of the amino acid boxes correspond to the coloring of the monomers in Fig. 1D and 1E.

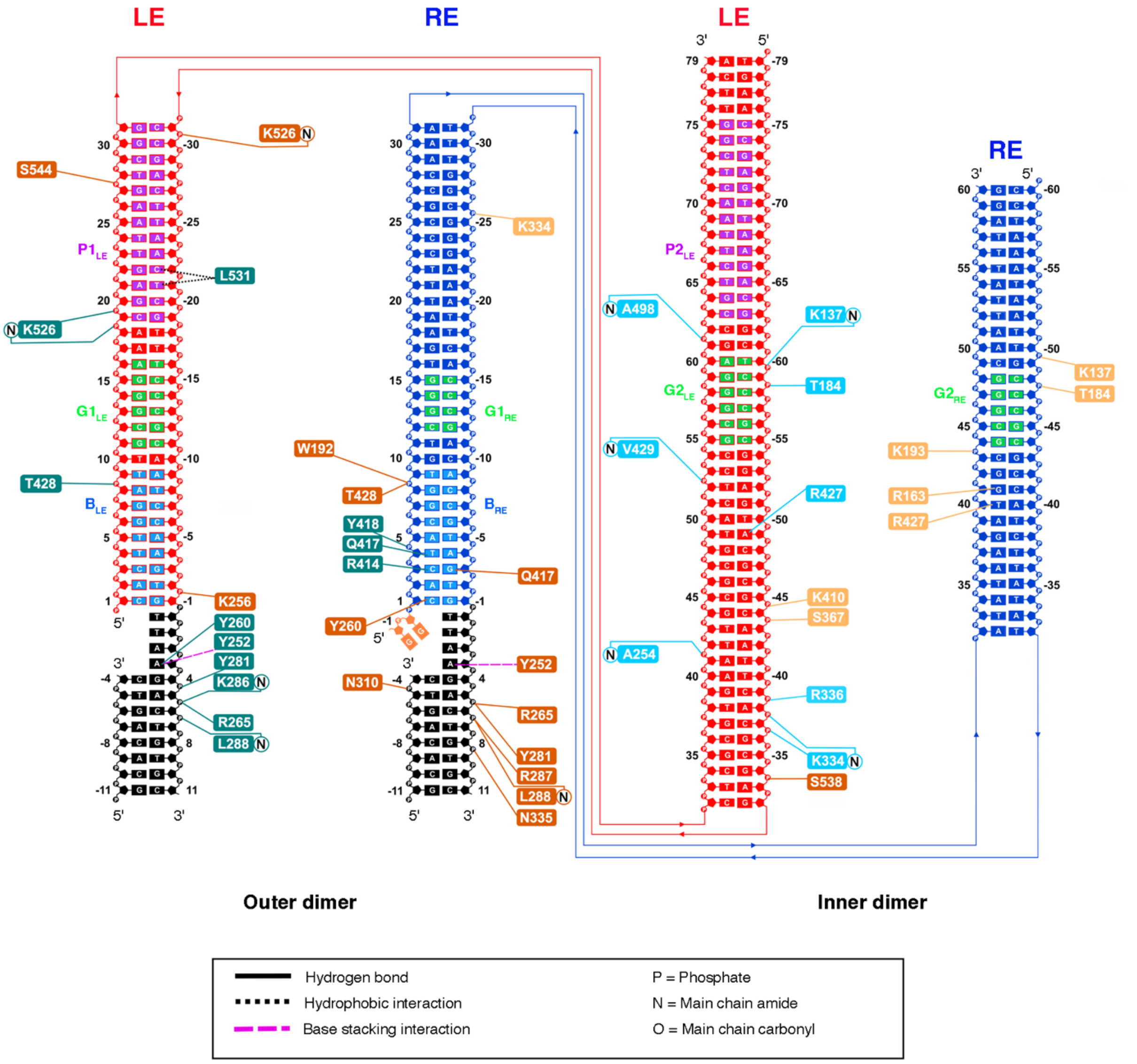


**Supplementary Figure 7. Schematic of the unique protein-DNA interactions present in outer and inner dimer of the pBat STC assembly.**

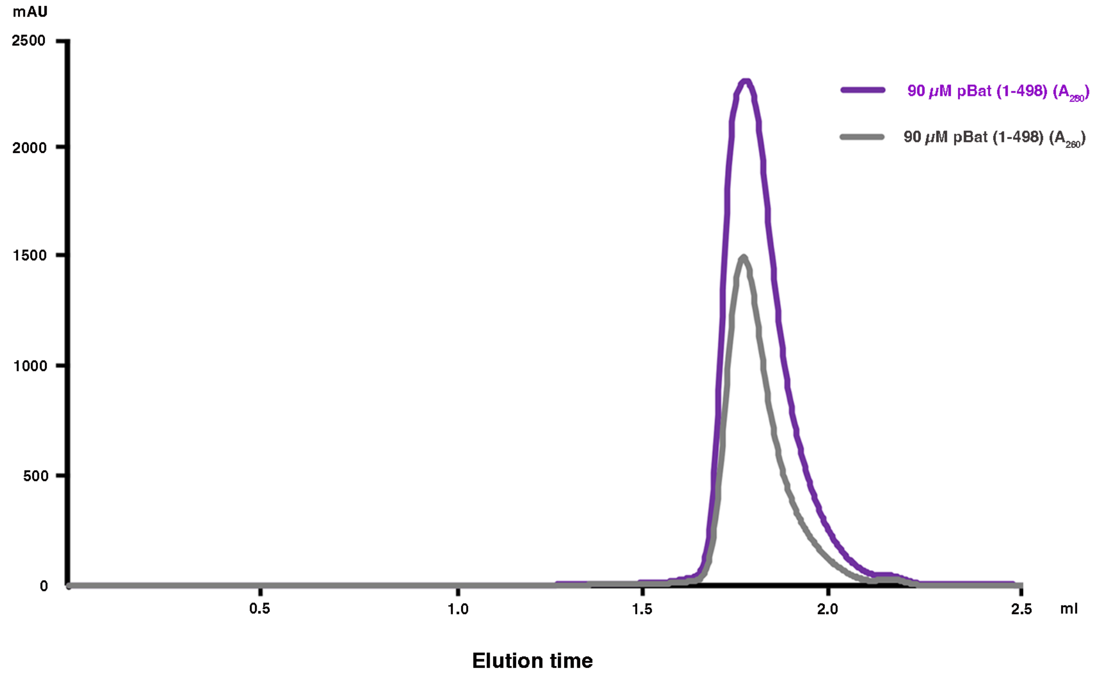


**Supplementary Figure 8. Elution profile of pBat (1-498) on a Superose 6 increase 3.2/300 column.**

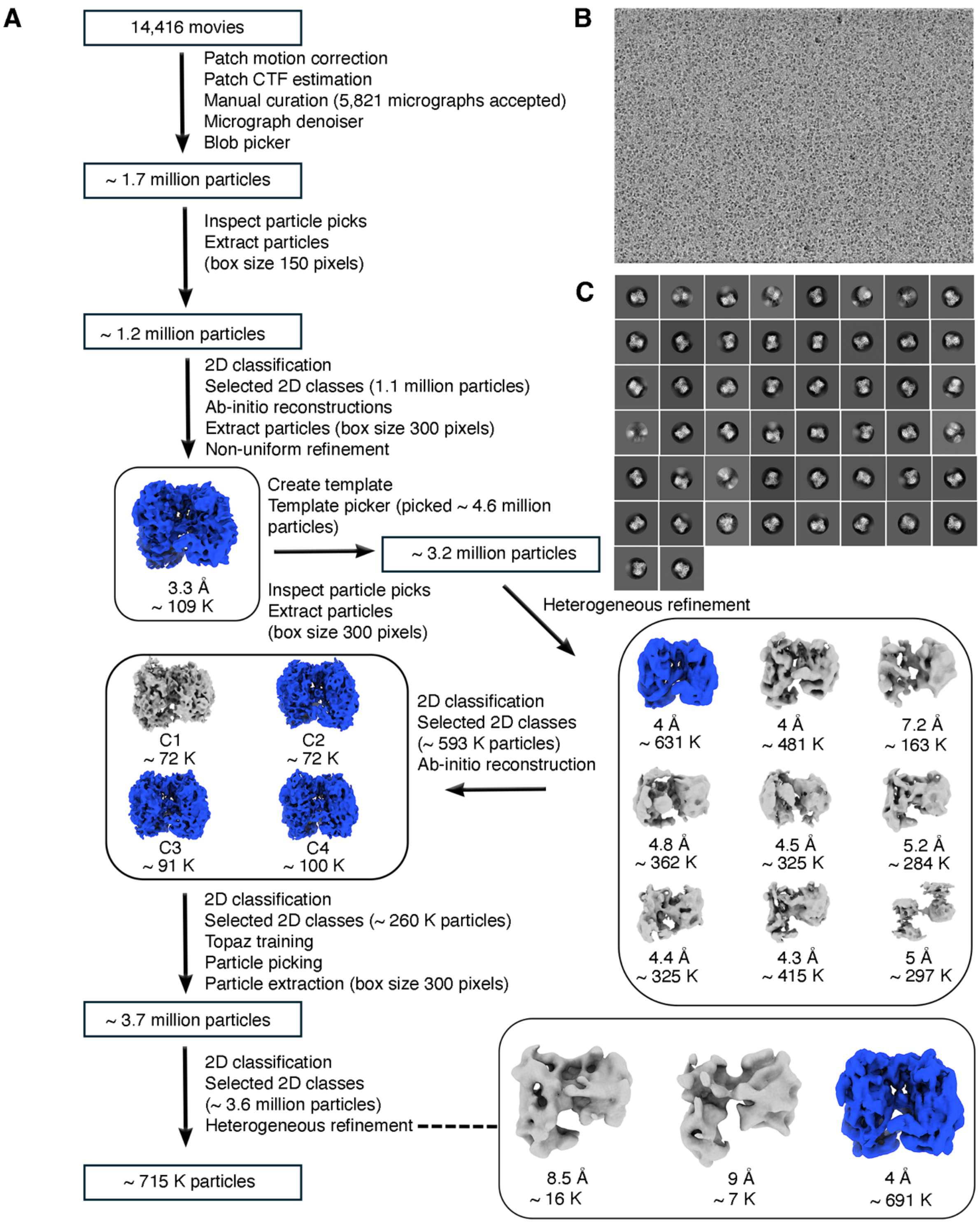


**Supplementary Figure 9. Cryo-EM data processing workflow of pBat (1-498)**. (A) Detailed workflow of cryo-EM data processing of pBat (1-498) in CryoSPARC. (B) A sample motion-corrected micrograph of pBat (1-498). (C) Reference-free 2D class averages of pBat (1-498) particles during the early stages of data processing in CryoSPARC.

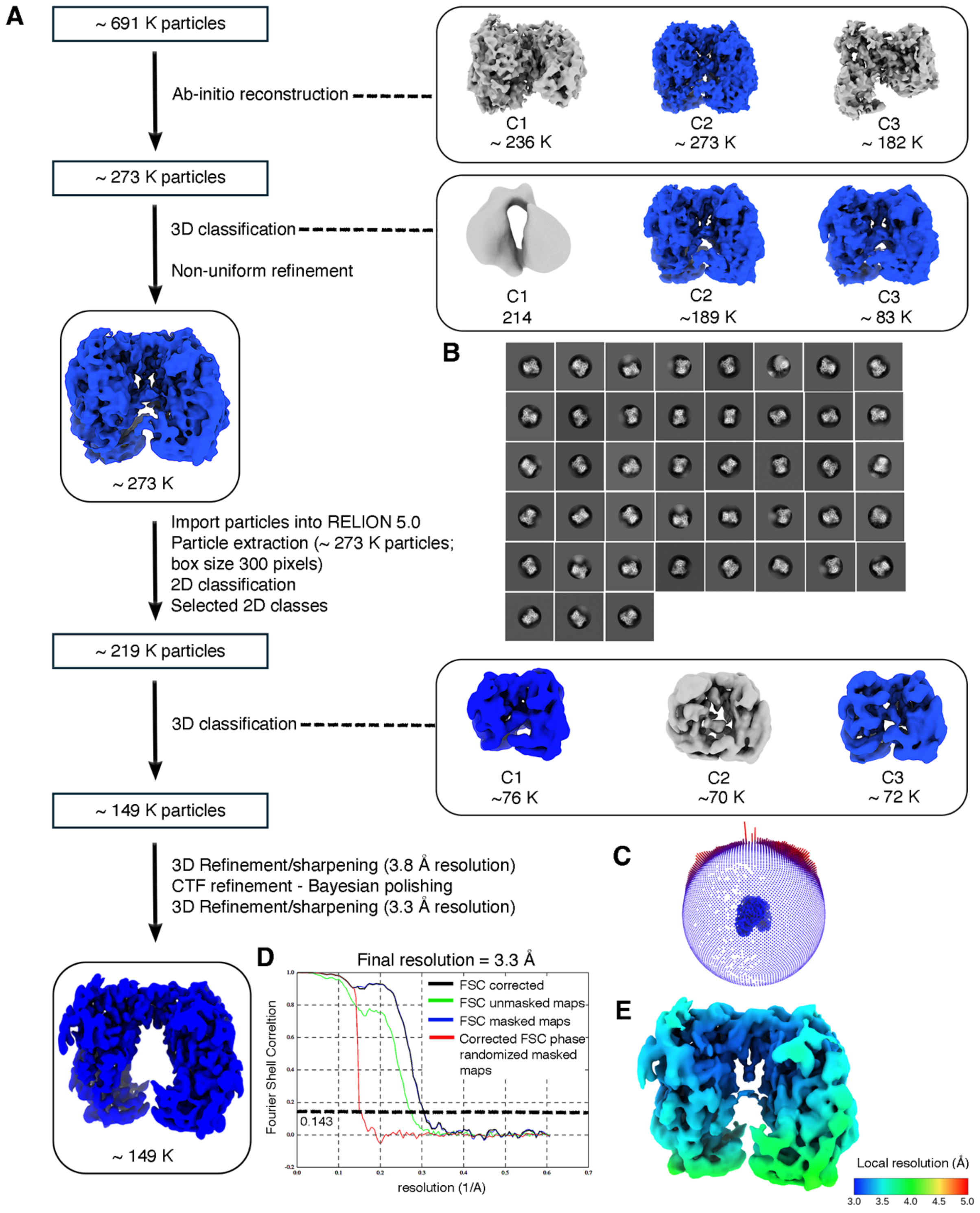


**Supplementary Figure 10. Cryo-EM data processing workflow of pBat (1-498).** (A) Detailed workflow of the cryo-EM data processing in both CryoSPARC and RELION. (B) The 2D class averages of pBat (1-498) particles in the later stage of data processing. (C) The distribution of particles view used for the final refinement. (D) The final resolution of the cryo-EM map was achieved at 3.3 Å at 0.143 FSC, although RELION overestimated the resolution due to preferred orientation. (E) Resolution distribution of the final reconstruction in RELION at a contour level of 0.005.

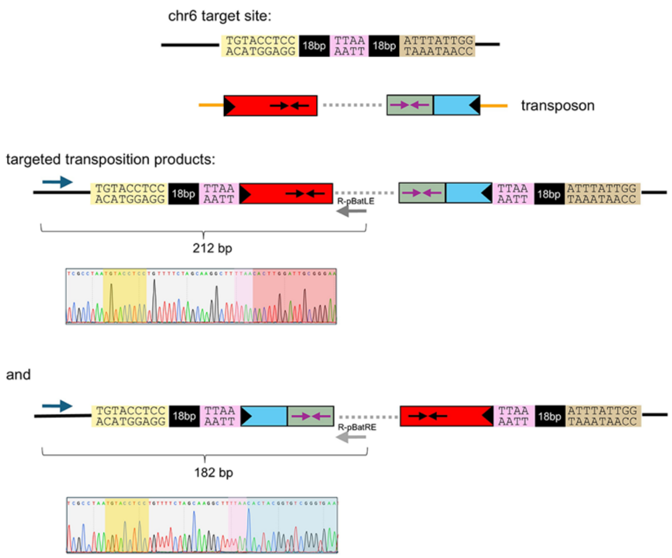


**Supplementary Figure 11. gPCR to detect integration in HEK 293T cells.** Schematic of the two possible products of transposon integration into the chr6 TALE target site, and examples of representative Sanger sequencing of gPCR products that confirm integration in HEK 293T cells.
